# Fluorescence Lifetime Imaging Microscopy (FLIM) visualizes internalization and biological impact of nanoplastics in live intestinal organoids

**DOI:** 10.1101/2025.01.27.635069

**Authors:** Irina A. Okkelman, Hang Zhou, Sergey M. Borisov, Angela C. Debruyne, Austin E. Y. T. Lefebvre, Marcelo Leomil Zoccoler, Linglong Chen, Bert Devriendt, Ruslan I. Dmitriev

**Author notes:** these authors contributed equally to this work (Irina A. Okkelman, Hang Zhou).

## Abstract

The increasing micro- and nanoplastic (MNP) pollution poses significant risks to human and animal health, yet the mechanisms of their accumulation and effects on absorptive tissues such as the gastrointestinal tract remain poorly understood. Addressing these knowledge gaps requires tractable models coupled to dynamic live cell imaging methods, to enable multi-parameter analysis at single cell resolution. Here we report a new method combining adult stem cell-derived small intestinal organoid cultures with multi-parameter live Fluorescence Lifetime Imaging Microscopy (FLIM) to study MNP interactions with gut epithelium. To facilitate this, we optimized live imaging of porcine and mouse small intestinal organoids with an ‘apical-out’ topology. Subsequently, we produced a set of pristine MNPs based on PMMA and PS (<200 nm, doped with deep-red fluorescent dye) exhibiting different surface charges, and evaluated their interaction with organoids displaying controlled epithelial polarity. We found that nanoparticles differently interacted with apical and basal membranes of the organoids and even showed a species-specific pattern of cellular uptake. Using a phasor-FLIM approach, we demonstrate better sensitivity of FLIM over conventional intensity-based microscopy. The ‘fluorescence lifetime barcoding’ enabled distinguishing different types of MNP and their interaction sites within organoids. Finally, we studied short (1 day)- and long (3 days)-term exposure effects of PMMA and PS-based MNPs on mitochondrial function, total energy budget and epithelial inflammation and found that even pristine MNPs could disrupt chemokine production and mitochondrial membrane potential in intestinal epithelial cells. The presented FLIM approach will advance the study of MNP toxicity, their biological impacts on gastrointestinal tissue and help tracing other types of fluorescent nanoparticles in live organoid and 3D *ex vivo* systems.

## Introduction

Plastics pervade every aspect of modern life, from packaging and household goods to applications in agriculture and medicine. Their durability, versatility and other attractive features have driven their widespread adoption but simultaneously contribute to their resistance to (bio)degradation. As a result, the use of slowly degrading polymers in everyday plastics resulted in the global accumulation of micro- and nanoplastics (MNPs) across diverse ecosystems, including soils, aquatic environments, the atmosphere, and even food sources like meat, drinking water, and vegetables. These MNPs interact with the biosphere in complex ways, giving rise to an emergent ecological niche termed the ‘plastisphere’^1–3^. Despite growing interdisciplinary efforts to mitigate plastic pollution through environmental remediation and the development of degradable polymers, the plastisphere is poised to remain a global societal challenge for decades to come.

Humans and animals are mainly exposed to MNPs via inhalation and ingestion. Disturbingly, MNPs have already been detected in human blood, the brain, the gut, the reproductive system and various other organs^4–8^. While precise estimates of MNP intake in livestock and other animal species remain unavailable, exposure levels are likely comparable. The presence of MNP in both humans and animals raises pressing questions about their potential health effects. Limited studies in rodents and human cell lines have shown that MNPs can negatively impact the intestinal epithelium, disrupting cell metabolism as well as triggering oxidative stress and inflammation in the gut^9–11^. However, it remains unclear whether these effects occur in human tissues or across diverse animal species in a physiologically relevant three-dimensional context. Real-world plastic particles can display a complex size and shape distribution and can be associated with contaminating metals, bio- and living matter traces. However, to establish a fundamental understanding of MNP uptake mechanisms and their effects on tissues, ‘model’, weathered or ‘pristine’ particles are needed^1,12^.

To address the complexity of the MNP interactions with tissues, the use of tractable 3D tissue models is essential. Advances in induced pluripotent stem cell and adult stem cell technologies now enable to recapitulate key aspects of tissue development and environmental interactions using organoid models. For instance, ‘mini-guts’ or intestinal organoids closely mimic the cellular composition, functionality, 3D architecture and extracellular matrix organization of the native intestinal epithelium, providing a robust platform to investigate MNP interaction with the gut epithelium and help producing multi-organ on-a-chip models^13–16^.

While organoid models are highly amenable to ‘omics’ and bulk analysis methodologies, their true complexity can be grasped best with imaging methods that provide live, multiparametric and quantitative readouts, together with long-term dynamic imaging, such as live cell fluorescence microscopy, full-field OCT and related approaches^15,17,18^. Fluorescence lifetime imaging microscopy (FLIM) presents such a method, allowing multiparametric and multi-dimensional 5D imaging (X,Y,Z, *t* and luminescence lifetime (tau, τ) dimensions)^18,19^. FLIM enables to study intra- and intercellular processes using biomarkers and a growing list of physical-chemical parameters pertinent to cell function, including inflammation and cell metabolism^18^. The latter is tightly connected with cell growth, differentiation, “live vs. death” decisions and the stem cell niche environment^20^, all of which can be replicated in 3D organoid models^15^. In addition to sensing and measuring cellular physiology in live organoids, FLIM can also be used to trace MNPs. The latter can exhibit unique and material-specific autofluorescence or can be loaded with various fluorescent dyes^18^. However, in respect to MNP, FLIM-based approaches focused so far only on the analysis of purified and separately present MNP in relatively simple sample solutions^21–23^. Such ‘intrinsic’ spectral properties and ‘mixing with the dye’ approaches can be challenged by the strong autofluorescence in live organoids^24,25^ and provide limited information on the MNP type, localization and composition.

To address the need for a more accurate analysis of the structure-activity relationships of MNPs, interacting with tissues, here we report a live cell FLIM approach, using optimized small intestinal organoid cultures and deep red dye-impregnated pristine nanoparticles for studying MNP interactions with the gut epithelium. We demonstrate that our approach enables efficient lifetime-based separation of different polymers and a highly sensitive MNP detection, which can be combined with long-term dynamic imaging of MNP internalization sites and their effects on cellular energy production mechanisms and inflammatory responses.

## Results and Discussion

### Design, spectral and fluorescence lifetime characterization of pristine nanoparticles mimicking MNPs

Various types of polymers can be used to produce dye-doped nanoparticles via nanoprecipitation and achieve controlled size and distribution^26–28^. Considering the need in live imaging, photostability, hydrophobicity and organoid live cell autofluorescence, we selected dibutoxy-aza-BODIPY dye (diBuO-aza-BODIPY^29^, exc. 700 nm/ em. 720 nm) as an emitter for doping the polymeric particles. For the ‘pristine’ polymer MNP and proof-of-principle method development, we selected negatively and positively charged polystyrene (PS) and poly(methylmethacrylate) (PMMA)-based structures^8,26,30^. Both PMMA and PS are abundant types of MNP^31^ and are widely used in bio- and nanosensing applications across the life science domains^8,32,33^. Using nanoprecipitation method, we produced 4 types of deep red emitting nanoparticles (NP A-D) with a comparable size and dispersity (100∼200 nm, based on dynamic light scattering, and 58∼134 nm, based on TEM, Fig. 1, S1 and Table 1). As expected, we found high brightness and deep red fluorescence (exc. 694∼703 nm with em. max. 717∼732 nm, depending on the polymer) with all 4 NP types (Fig. 1B, Table 1). Interestingly, depending on the polymer and the presence of the charged groups, the fluorescence lifetimes (analyzed by three individual methods, see Table 1 and Fig. 1C) in water for all the 4 types of the NPs showed a broad range from 1 to 4 ns with the multi-exponential fluorescence decays. The fluorescence lifetime of the dye depends on the chemical structure of the polymer and observed differences could originate, for example, from partial aggregation of the dye in the matrix or the sensitivity of the luminescent properties to different environmental parameters^27,34^. For NP A, consisting mostly of PMMA and acrylate units bearing quaternary ammonium groups, we observed a strong influence of the environment on the fluorescence lifetime (roughly estimated by τ_ϕ_ phasor lifetimes), generally decreasing from 3.7 in deionized water to 2.7 and 1.5 ns (Fig. 1D) in presence of FBS and in contact with live organoids, respectively. In contrast, NP D (Fig. 1E), that are based on copolymer of styrene and maleic acid, showed generally constant fluorescence lifetime: τ_ϕ_ =1.96 ns in water, imaging medium, 25% FBS or when taken up by organoids. Such striking differences in fluorescence lifetimes, based on the type of the polymer structure and charge represent an attractive ‘barcoding’ feature to distinguish between different types of MNP within the same spectral (excitation-emission) channel. Importantly, τ_ϕ_ and τ values observed on different detection platforms matched well, confirming reliability of the FLIM measurements (Tab. 1). We subsequently illustrated this ‘barcoding’ feature by fusing drops of NP A and B solutions (NPs with the longest and the shortest fluorescence lifetime in water) and performing phasor FLIM analysis (Fig. S2). The resulting phasor cluster, produced from individual points with G and S coordinates counted from fluorescence decays of corresponding FLIM image pixels, displayed a typical indicative elongated shape and was aligned between the positions of the two distinct NP A and NP B emitting species clusters (Fig. S2A-B). According to the phasor principle the distance of the fused cluster points from one of the distinct species clusters reflects the weight of this species in a mixture at the corresponding pixel location on the image^35^. Thus, the analysis of fused phasor cluster point coordinates can be applied for detecting the multiple emitting species with spatial resolution, if individual fluorescent species lifetimes are not strongly affected by the local environment. With the assumption that fluorescence lifetime is an indicative parameter of distinct emitting species (including system noise), a simple detection of the phasor cluster points with the corresponding G and S coordinates can be used to detect the potentially rare events of NP uptake by tissues.

**Figure 1:**
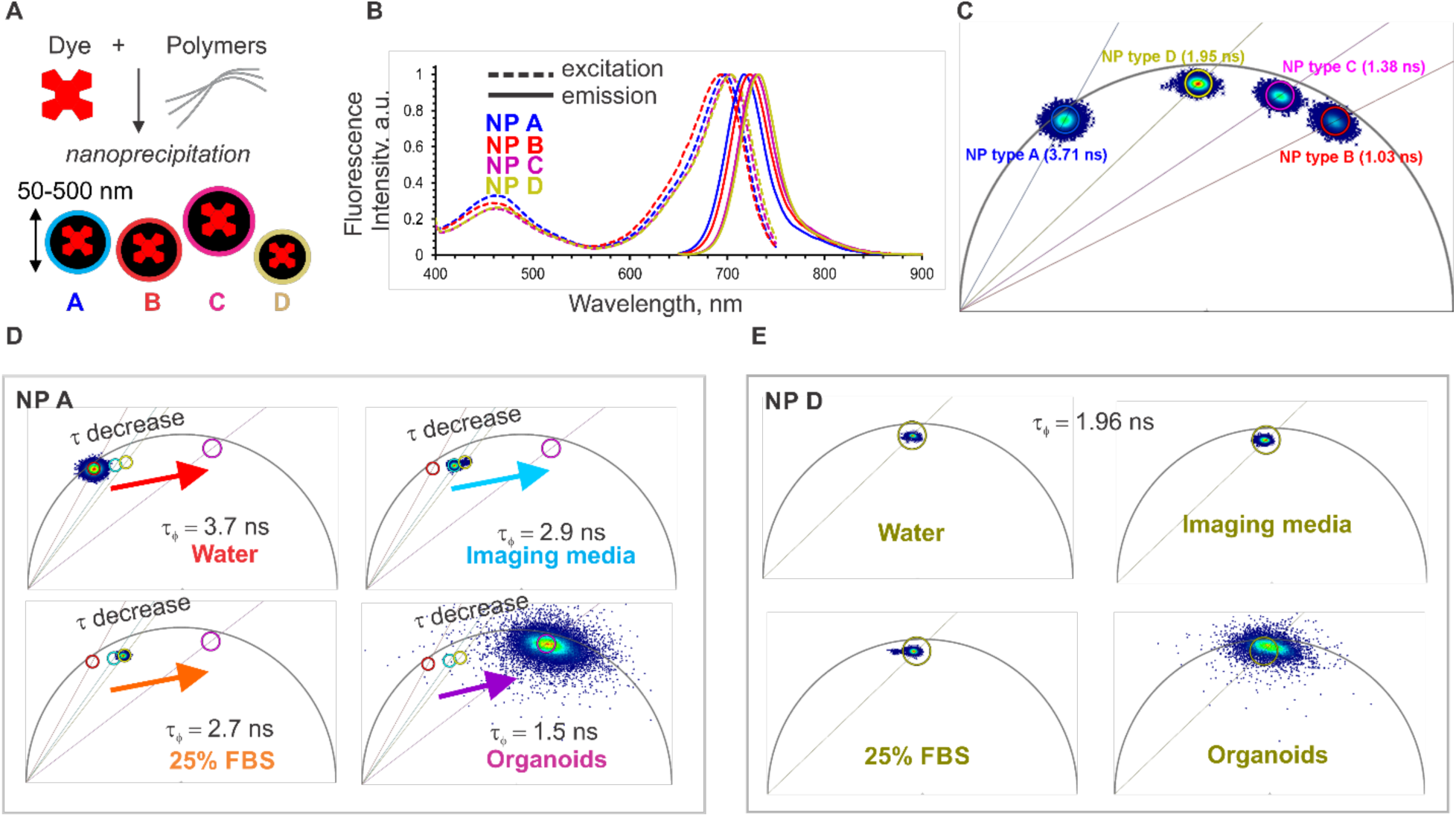
Design, spectral and fluorescence lifetime characteristics of the model nanoplastics NP. **A-D. A.** Schematics of NP design. **B.** Normalized fluorescence excitation and emission spectra of NP A-D. **C.** Phasor-FLIM characterization of NP A-D (0.5 mg/ mL, deionized water, 37 °C). Centre τ_ϕ_ values indicated in brackets. **D, E:** Effect of the environment on the phasor plots of NP A (D) and NP D (E), respectively. Nanoparticles were measured in all solutions at 0.5 mg / mL, with exception of organoids (10 µg/mL, incubated for 18-24 h).

**Table 1.** Properties of dibutoxy-aza-BODIPY-doped nanoparticles. Z_av_ values obtained via dynamic light scattering (nanoparticles in deionized water) and TEM are presented. τ values were measured on fluorescence spectrometer (‘Fluo’), τ_ϕ_ and intensity-weighed τ_m_ values were obtained on FLIM microscope (‘FLIM’) via phasor or decay fitting approaches (see Methods for more details).

| NP type (polymer) | $I_{exc}$, nm | $I_{em}$, nm | $Z_{av}$, nm (DLS) | zeta-potential, mV (DLS) | $Z_{av}$, nm (TEM) | $\tau$, ns (Fluo) | $\tau_\phi$ in water ('FLIM', phasor), ns | Intensity weighted $\tau_m$ in water, ns (FLIM, decay fitting) | $\chi^2$ |
| --- | --- | --- | --- | --- | --- | --- | --- | --- | --- |
| A (RL100) | 698 | 717 | n.d. | n.d | 78 | 4.4 | 3.71 | 4.02 | 2.447 |
| B (PMMA-MA) | 694 | 723 | 99 | -29 | 81 | 1.1 | 1.03 | 1.15 | 0.938 |
| C (PS-PVP) | 703 | 729 | 200 | -7 | 99 | 1.5 | 1.38 | 1.53 | 1.145 |
| D (PS-MA) | 703 | 732 | 166 | -28 | 135 | 2.3 | 1.95 | 2.21 | 1.216 |

To estimate the reliability of the dibutoxy-aza-BODIPY-labeled NP cluster points position on a phasor plot in comparison to the system noise signal (which includes optical noise, short noise or thermal noise), we reconstructed phasor clusters for TCSPC-FLIM measurements of different dilutions of NP D in water (0.58-500 µg/mL) and compared them to the signal from water containing no NPs, which can be referred as a “system noise” sample (Fig. S3). The dilution of homogeneous NP D solution led to a decrease in fluorescence intensity and accordingly to a decrease in the photon rate per pixel. This decrease in the pixel photon count upon ∼1000-fold dilution (from total photons count 9.9*10^7^ for 500 μg/mL solution decay to 1.4*10^5^ photons for 0.58 μg/ mL solution decay measured for 512×512 pixel resolution image, binning 1) correlated with the increase in NP D phasor cluster points dispersion from the theoretical center of the cluster, indicated as the center of the circular ROI mask (Fig. S3, magenta). However, the principal position of the NP D phasor cluster on a plot remained intact within the same ROI mask (magenta ROI, Fig. S3). Importantly, we could narrow down the scattering by applying the pixel binning and improving the photon counts per pixel. At the same time, we did not observe any random localization of the noise cluster points on a phasor plot within the area of the NP D phasor ROI. In contrast, pixel binning reduced noise pixels scattering by clearly segregating their localization in a short lifetime zone of the phasor plot (yellow ROI). Thus, even with a considerable scattering of the points from the main cluster position, the dibutoxy-aza-BODIPY-labeled NP cluster could be reliably detected with the pixel binning approach, improving the event detection reliability. Remarkably, 0.58 μg/mL NP D concentration in water still was not a limit for the detection of the NP D cluster on a plot without shifting its general position towards the noise and corresponding changes of the τ_φ._ The presented data is in contrast to a similar approach^21^ utilizing TCSPC-based phasor plot analysis of NP in water, where due to low signal-to-noise ratio the achieved limit of detection of non-stained polystyrene nanoparticles was only 10 μg/mL. This difference in sensitivity might be due to a different microscopy instrument set up as well as the NP labeling with a near-infrared aza-BODIPY dye, which is characterized by a fairly high quantum yield^36^ and allows to keep a high signal-to-noise ratio upon dilution of the NP.

### Visualization of apical-out organoid polarity reversion via live microscopy

An ‘apical-in’ or Basal-Out (referred as BO in the text) topology of small intestinal organoids represents a major limitation of this model for studying direct interactions of the apical membrane with nutrients, microbiota and pathogens. While this has been addressed by microinjection or partial disaggregation to crypts and re-assembly^15^, a more elegant approach was introduced by the group of Amieva^37,38^ for human enteroids, via depletion of ECM and subsequent transition to suspension culture. We subsequently adapted this protocol to the cultures of porcine and mouse small intestinal organoids and confirmed the phenomenon of the polarity reversion, using staining of F-actin as a conventional marker of the apical membrane^38^ (Fig. S4). Since the process of polarity reversion can differ from organoid to organoid, requiring live monitoring of organoids topology, we evaluated two tracers enabling labeling of an Apical-Out (AO) topology in live small intestinal organoids: fluorescent wheat germ agglutinin (WGA) and Nile Red, labeling the cell membrane and lipid droplets (LD), respectively (Fig. S5, Fig. 2). Both tracers display attractive spectral and fluorescence lifetime properties (∼2.1 ns for WGA-Alexa Fluor 488 and ∼3.3 ns for yellow-red emission of the Nile Red (Fig. S5B, S5C, S5E, Table 2).

**Figure 2.**
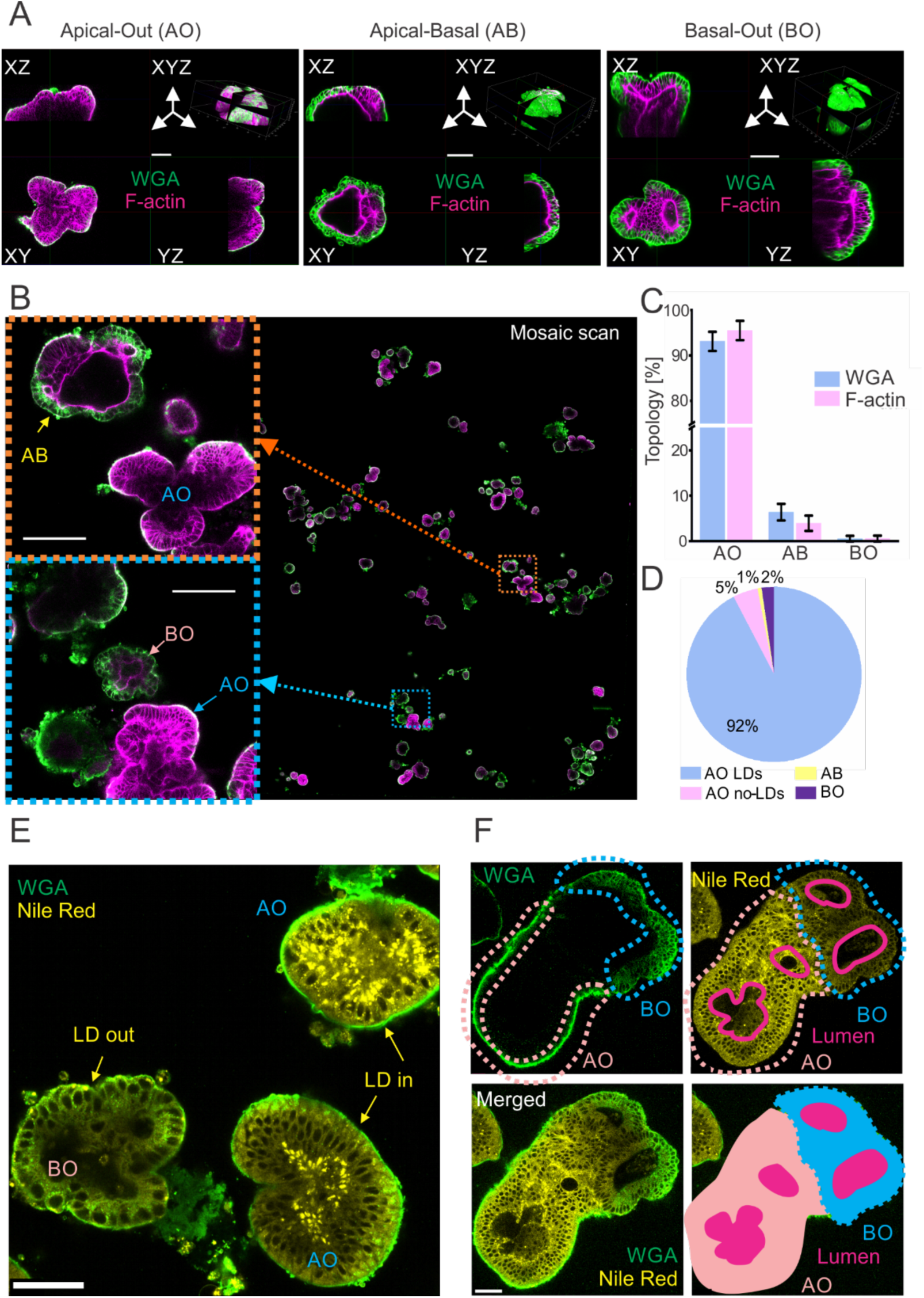
Fluorescent Wheat Germ Agglutinin (WGA) conjugate and Nile Red enable validation of the apical-out topology in live small intestinal organoids. **A:** Representative 3D confocal images of WGA-Alexa Fluor 488 and phalloidin–Texas Red (F-actin labeling) co-stained organoids with apical-out (AO), apical-basal (AB) and basal-out (BO) topology. Scale bar is 100 µm. **B:** Large field mosaic scan of PFA-fixed pig intestinal organoids after 18 h of polarity reversion, co-stained with fluorescent WGA and phalloidin. *Left:* representative images of AO, AB and BO organoids, indicated on mosaic image (*right*). Scale bar is 100 µm. **C:** Comparison of the yield of AO, AB and BO topology, estimated with WGA staining and F-actin labeling, respectively (AO organoids: 93.1% / 95.5%, BO organoids 0.53% / 0.55%, analyzed from 4 mosaic scanned images, see table ST3). **D:** Total quantification of topology, observed with WGA and Nile Red labeling for polarity reverted organoids: AO with lipid droplets (AO LDs) and AO without lipid droplets (AO no-LDs), AB and BO, quantified from mosaic scanned image. **E:** Lipid droplets display a characteristic distribution in AO organoids, contrasting with BO. **F:** Co-staining with WGA and Nile Red reveals organoid structure in apical-basal organoid (AB). Scale bar is 50 µm.

**Table 2.** Fluorescent probes used in live microscopy of organoids. Fluorescence lifetime was measured using FLIM microscope (see ‘Microscopy’).

| Probe | Physiological parameter | Stock concentration (Solvent) | Staining concentration | Excitation / Emission (nm) | Fluorescence Lifetime ($\tau$, ns), [citation] |
| --- | --- | --- | --- | --- | --- |
| AF488-WGA | Binds N-acetyl-D-glucosamine and sialic acid; mucous membranes<sup>70</sup> | 1 mg/mL (PBS) | 4 μg/mL | 490/495-520 | 1.2-2.6 ns [this study] |
| Fluorescein-UAE-1 | Glycoproteins, glycolipids with α-fucose residues<sup>71</sup> | 5 mg/mL (PBS) | 5 μg/mL | 490/495-520 | NA (No staining observed) |
| Nile Red | Solvatochromic dye. Lipids, LDs, cytoplasm. | 1 mM (DMSO) | 2 μM | 490/588 - 620 | 2-4 ns<sup>69</sup> |
| Mito Tracker Green (MTG) | Mitochondria | 10 μM | 100 nM | 490/497 - 551 | 0.5-1.3 ns [this study] |
| Mito Tracker Red (MTR) | Mitochondria | 10 μM | 100 nM | 581/594 - 654 | 0.8-1.2 ns [this study] |
| TMRM | Mitochondrial polarization | 10 μM (DMSO) | 10 nM | 540/559 - 585 | 1.5-2.5 ns<sup>52</sup> |
| Phalloidin-Texas Red | F-actin labeling in fixed cells | 66 μM | 0.132 μM | 591/588 - 625 | 3-3.3 ns [this study] |

WGA: wheat germ agglutinin, UAE-1: Ulex europeaus agglutinin- 1; TMRM: tetramethylrhodamine methyl ester;

To evaluate WGA labeling of the organoids, we first performed a global analysis of the small intestinal organoid topology after 18 h suspension culture (Table S1) and found 3 topological states: Apical-Out (AO, mainly outer membrane labeling with WGA, Fig. S5A and S5G), Basal-Out (BO, deeper cellular borders labeling with WGA and internalization of WGA, Fig. S5D and S5G) and Apical-Basal (AB, both AO and BO characteristic staining visible in different parts of the organoid)(Fig. S5G). We then validated organoid topologies with co-localization experiments using live WGA staining followed by fixation and F-actin staining (Fig. 2A). To quantify the total number of each type, we performed mosaic imaging of organoids co-stained with WGA and F-actin (Fig. 2B) and found a similar percentage of AO (WGA: 93.1%, F-actin: 95.5% and BO (WGA: 0.53%, F-actin: 0.55%) topology in organoids with both labelling approaches (Fig. 2C).

Interestingly, Nile Red (yellow color) provided similar topological information on organoids through the spatial distribution of lipid droplets (LDs), localized at the basal membrane side^39^, i.e. within the internal region of AO and external region of BO organoids (Fig. 2E). To confirm the lipid droplet distribution as a reliable marker for AO identification, we classified organoids to AO, AB and BO by WGA labeling and then quantified the lipid droplets in AO organoids via mosaic imaging (Fig. S5H). We found that most AO organoids contained LDs (AO with LDs: 92%, AO without LDs: 5%), confirming Nile Red-labeled LDs as a reliable identifier for AO morphology (Fig. 2D). Furthermore, we observed that the Nile Red signal was higher in AO compared to BO organoids, while WGA staining intensity remained similar between these two groups (Fig S5D, S5E). Statistical analysis within defined AO region of interest (ROI) and BO ROI from mosaic scanned images using FIJI (Fig. S5F, S5H), confirmed Nile Red intensity being a reliable marker for discriminating between AO and BO organoids (p=0.006).

Collectively, combining Nile Red with WGA-Alexa Fluor 488 provided better information on the organoid topology in live organoid imaging. Importantly, Nile Red is compatible only with live imaging as it does not remain in organoids after fixation, in contrast to WGA labeling. An advantage of Nile Red and WGA co-staining is in deeper information on the internal organoid structure, allowing identification of lumen (for BO structures) and pseudo-lumen regions (for AO structures) within the live organoids (Fig. 2F). This is in contrast with many other fluorescent tracers, such as UEA-1, live actin tracking probes (e.g. CellMask Actin Tracking Stains and SPY-actins, data not shown) and mitochondrial stains (MTG, TMRM and MTR), which demonstrated negative or differential staining within the population of organoids in suspension or parts of the individual organoids (Fig. S5I). This phenomenon can be explained by a potential presence of mucus in the AO organoids or a preferential uptake of the dyes through basal or apical membranes of the gut organoids, which has to be further elucidated.

### Nanoplastics display specific accumulation in small intestinal organoids

Using our optimized polarity reversion and live FLIM methodology, we exposed live organoids to the 4 types of NP (A-D) (Table 1, Fig. 3A). With the ‘standard’ incubation time (18-24 h) and staining concentration (10 μg/mL), routinely used in stainings with fluorescent and phosphorescent nanosensors^25,27,28,34,40–42^, we observed differences in the interaction of NP with porcine intestinal organoids: NP C (PS-PVP) showed no uptake, NP A (PMMA with quaternary ammonium groups) mostly accumulated at the organoid membrane, while both negatively charged NP types B (PMMA-MA) and D (PS-MA) showed specific intracellular uptake (Fig. 3B). Phasor plots also revealed an appearance of recognizable clusters within the specific lifetime distribution zones (see Fig. 1C) for NP A, B and D after NP exposure, while NP C and control (no NP) had no detectable signal (Fig. 3B). Thus, NP A, B and D demonstrated interaction with and internalization by intestinal epithelial cells, independently of the backbone structure (PMMA or PS) and the presence of charged groups (quaternary ammonium or methacrylate groups). This pattern was consistent across 3 different small intestinal organoid lines (porgJ-2, Fig.3; porgJ-3 and porgJ-4, Fig. S6) developed from 3 animals, suggesting animal-independent general uptake of these NPs by the pig intestinal epithelium.

**Figure 3.**
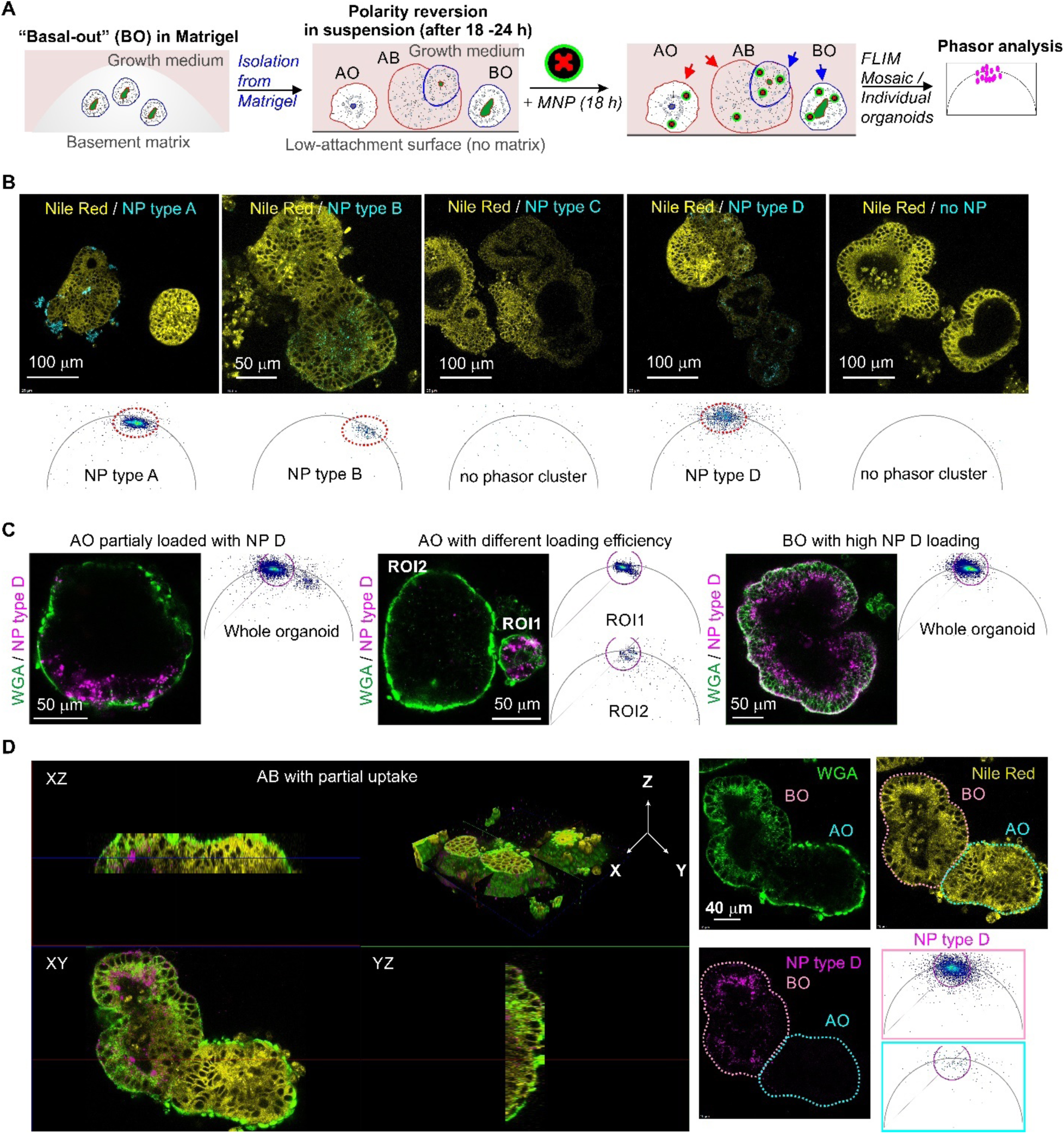
MNPs display diverse staining and uptake in the small intestinal organoids. **A:** Scheme of experimental workflow. **B:** Representative confocal fluorescence images and corresponding phasor plots of the pig small intestinal organoid incubated with NP A-D (10 µg/ mL, 18 h), co-stained with Nile Red. NP A, B and D displayed signals on both fluorescent intensity images and phasor plots, whereas type C showed no signal, similar to control. **C:** Different types of uptake of NP type D uptake (10 µg/mL NPD, 20% laser power) into organoids, co-stained with WGA, with respect to the topology and size. *Left:* partial NP D uptake in AO, showing high pixel signal on a phasor plot. *Middle:* organoid size-dependent NP D uptake in AO. ROI 1: enhanced NP D pixel signal in a small AO, ROI 2: reduced NP D pixel signal in larger organoid. *Right:* Homogeneous NP D distribution in BO with strong signal on a phasor plot. **D:** Topology-dependent uptake of NP D (magenta) in apical-basal organoid (AB), co-stained with WGA (green) and Nile Red (yellow). *Left:* 3D reconstruction shows distinct NP D distribution in AO and BO regions. *Right:* representative fluorescence image of AB organoid taken from 3D reconstruction. AO and BO regions defined by WGA and Nile Red signals, with corresponding NP type D phasor FLIM plots of these regions (bottom right).

Following WGA labeling, we found that NP D uptake depended on the organoid topology: thus AO organoids exhibited both high (ROI1, Fig. 3C middle panel) and low efficiency loading patterns (ROI2, Fig. 3C middle panel), in some cases belonging to the same organoid body (AO organoid with partial loading, Fig. 3C left). At first glance most AO organoids displayed lower NP D internalization in contrast to BO organoids (Fig. 3C, left panel) or BO zones in AB organoids, showing high efficiency and uniform uptake of NP D. This was confirmed by the live 3D reconstruction of AB organoid co-labeled with WGA and Nile Red (Fig. 3D). Differences in uptake from either apical or basal membranes imply potentially different routes of MNP delivery into the cells *in vivo*: either from the lumen of the gastrointestinal tract (e.g. with food or water) or from the bloodstream, i.e. with the injections.

We reasoned that such differences in interaction with cell membranes, caused by the type of MNP and the composition of the cell membrane, could also result in species-specific differences in MNP uptake. To test this, we produced AO intestinal organoids from Lgr5-GFP reporter mice^43^ (Fig. S7). Interestingly, we rarely observed BO and AB mouse organoids in the analyzed organoid population. Thus, we focused only on the interaction of NPs with AO organoids. The AO topology was confirmed by assessing the lipid droplet staining distribution with Nile Red staining and the shifted position of nuclei towards the basal membrane in polarity-reverted organoids^37^ (Fig. S7). In addition to A, B and D types of NP, we also detected rare events of NP type C accumulation in mouse AO organoids (Fig. S7C). For NP C, two different loading patterns were found: (i) low intensity zones (magenta mask, Fig. S7) within the organoid cells with lifetime ∼1.35 ns (similar to the fingerprint lifetime of NP C in water, Fig. 1C) and bright particles in rare cells with a lifetime ∼ 1.95 ns (red mask, Fig. S7D). This contrasted to the control group of unloaded organoids, where even the combined phasor plot from 8 individual microscopy images did not show appearance of the similar phasor clusters (Fig. S7E). Overall, all tested NP types tended to accumulate in specific cell zones (GFP-negative differentiated cells and ‘low intensity GFP’ early differentiated daughter cells) in organoids, with almost no accumulation within Lgr5-GFP^high^ intestinal epithelial stem cells. The exception was found only for NP A (Fig. S7A), where ROI-based phasor plot analysis revealed accumulation inside stem cell zones. In contrast to a predominantly surface membrane clustering of NP A in pig organoids, in mouse organiods NP A accumulated only inside the cells. Collectively, these data highlight the importance of species-specific effects of MNP interaction with absorptive intestinal epithelium and prompt for further investigation using controlled cell and mucus composition and the growth medium, which can affect cell metabolism and senescence^44,45^.

Finally, we studied the rate of MNP uptake in pig jejunum organoids by focusing on NP type D (Fig. S8). Using a confocal microscopy with mosaic scanning we continuously monitored organoids for 24 h after addition of NP D. Strikingly, NP D adhered to the membrane of some individual organoids already after 30 min (Fig. S8A). Subsequently, the MNP displayed slow internalization with rare particles detected inside organoids after 5 h of incubation (Fig. S8B) with almost complete uptake (low signal on the organoid surface) after 24 h (Fig. S8C). This suggests that in real-life situations, MNP may require a short exposure in order to ‘stick’ to the epithelial cells, while their physiological effect can occur with delay, caused by cell-specific internalization and accumulation.

### Phasor FLIM event counting approach reflects the number of MNP internalization events

As mentioned above, AO organoids displayed generally lower nanoparticle accumulation than BO organoids. We therefore decided to have a closer look on NP D accumulation in the pig polarity reverted organoids in respect of their apical-basal topology. To increase the number of MNP-loaded organoids we performed NP D loading during the polarity reversion process. Indeed, we observed a similar loading tendency in the organoids, where NP D internalization in BO or AB organoids was the highest with particles concentrating at the basal membrane and some localized at the apical membrane (Fig. 4 A,B). In contrast to BO organoids, which always displayed NP D uptake, AO organoids showed two distinct patterns: (i) organoids with MNP localized in vesicle-like NP bodies inside cells (Fig. 4A bottom panel) and (ii) organoids displaying no MNP accumulation (Fig. 4B bottom panel). NP D accumulation inside organoids in all positive cases was clearly recognizable by the appearance of characteristic NP D phasor clusters on the phasor plots reconstructed from the organoid ROI. We noticed that the number of phasor cluster points closely reflected the amount of fluorescent NP D bodies in the organoids. Knowing that every plotted phasor point corresponds to a pixel in the FLIM image with a certain photon count data, we hypothesized that simple point counting in phasor clusters from the characteristic lifetime zone would reflect the total number of NP bodies inside organoids. Thus, we evaluated fluorescence lifetime events detection to count internalized MNP.

**Figure 4.**
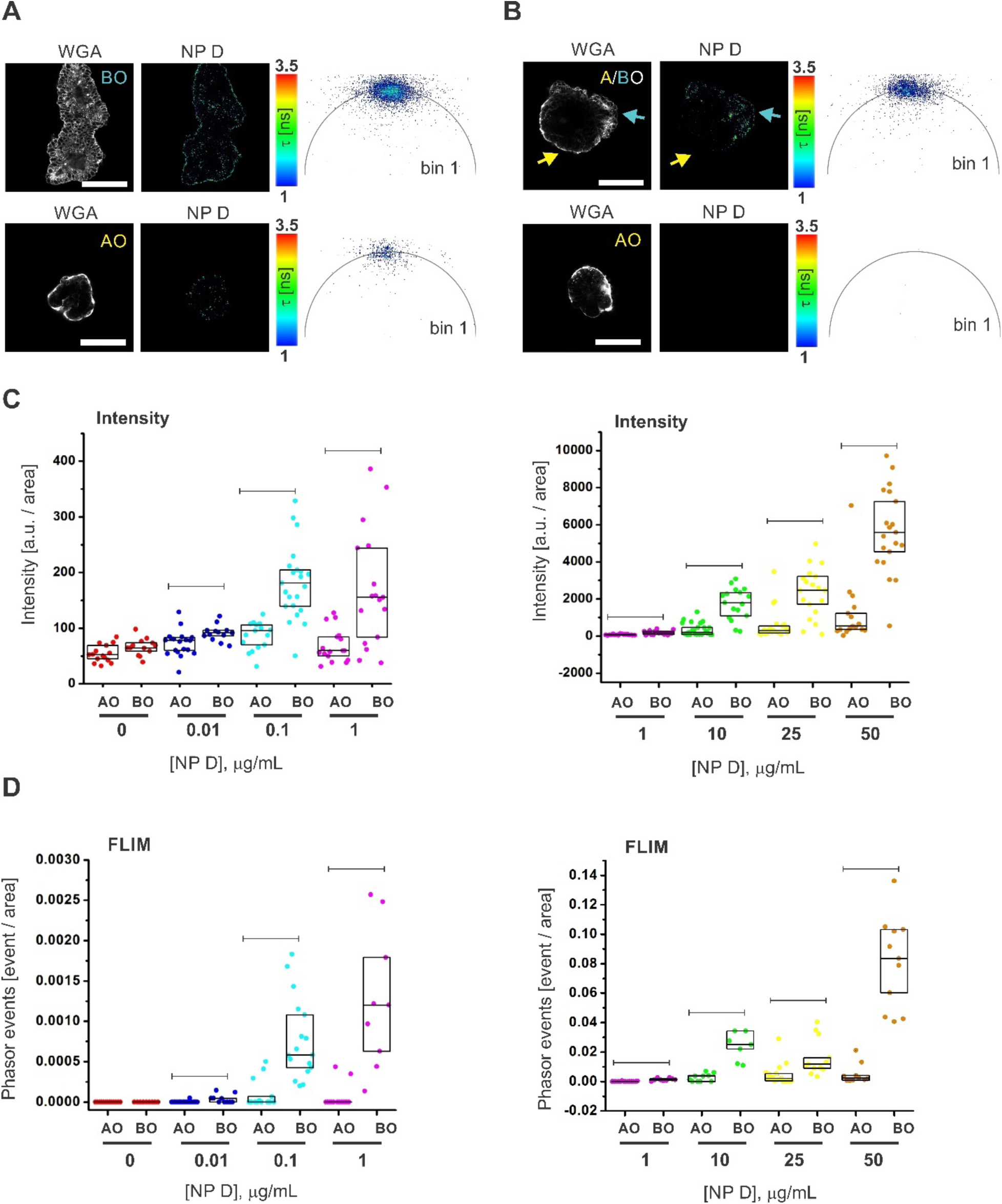
Phasor FLIM event counting approach estimates MNP uptake in organoids with improved reliability over a broad concentration range in comparison to intensity-based detection. **A, B:** Representative images of NP D uptake (10 µg/mL, 18 h) in pig intestinal organoids in respect to their apical-basal topology with high loading in both AO and BO (A) and high loading in BO and zero at AO (B). Scale bar is 50 µm. **C, D:** Comparison of NP D uptake in AO vs. BO organoids as a function of loading concentration (0-1 µg/mL, left panel and 1-50 µg/mL, right panel) on a widefield fluorescence microscope (intensity-based approach, C), and confocal FLIM microscope (phasor FLIM event counting approach, D). FLIM events were counted from the phasor plots reconstructed in the napari phasor plugin from the exported list of G and S coordinates. Results of one of the two independent experimental replicates are shown. Both C and D data were produced from the same samples. The box charts represent 25, median and 75 percentiles with dots corresponding to individual intensity or event count square ROI square ROI normalized values. Attribution of organoids to AO and BO-topology groups was done based on WGA staining. BO group also includes AB organoids. Statistical comparison between AO and BO groups over a range of loading concentrations was performed by Mann-Whitney test (lines represent detected statistical difference, p <0.05).

We also tested if the pixel binning could improve phasor clustering for NP accumulated in organoids, similar to the solution-based experiments with NP (Fig. S3). To have a more dispersed phasor cluster of NP D, we decreased the NP concentration to 1 µg/mL and performed imaging of non-loaded and NP D-loaded BO organoids with different excitation power (10 and 30 %), keeping all other imaging acquisition parameters unchanged. Phasor plots were reconstructed from the imaging data sets by variation of pixel binning (1, 3 and 6) and phasor thresholds (1 and 4) in LAS X software. No random points or cluster were detected in the lifetime zone of τ_ϕ_ ∼2-1.96 ns (Fig. S9, magenta circular ROI) on any of the reconstructed phasor plots of control organoid image when varying the excitation intensity. Application of a threshold parameter reduced the amount of noise cluster points, while pixel binning clustered points in a short lifetime zone within corresponding circular ROI (Fig. S9, yellow ROI). In contrast, a dispersed cluster of points appeared on the phasor plots (bin 1) of the NP-loaded organoids. The number of points increased by changing the excitation intensity to 30%. Some cluster points had a distinct drift toward the “noise” ROI, which was mitigated by exclusion of the noise affected pixels with the threshold application. Application of pixel binning further decreased the dispersion, while keeping the cluster position within the same lifetime zone. The minor shift of the cluster toward the longer lifetime zone could be explained by the improved photon statistics of individual pixels, making the phasor points position less sensitive to the noise impact during the calculation of the G and S coordinates. Overall, our observations consolidated the approach of implementing pixel binning and setting thresholds in addition to median filter-based point grouping for a robust reconstruction of phasor clusters. Thus, we confirmed the advantage of pixel binning^46^, efficiently improving a signal-to-noise ratio and making detection of a one-point event more reliable in case of MNP.

Next, we compared the phasor FLIM event counting approach with conventional intensity-based fluorescence measurement of NP D uptake in organoids over a range of loading concentrations 0-100 µg/mL. Examples of reconstructed phasor plots of organoids loaded with different NP D concentrations are shown in Fig. S10. First, we organized intensity and phasor FLIM events data based on the organoid topology and NP loading concentrations and compared values of AO and BO organoids in each ‘concentration’ group. Both intensity and event number values were higher in BO organoids as compared to AO organoids across all loading concentrations (0.01 – 100 μg/mL), except for the control group (Fig. 4 C, D; Table S3, 4). This result supports our observation (Fig. 4A) and confirms that both analysis approaches similarly reflect NP uptake in organoids.

Seeing that AO and BO organoids demonstrated a completely different NP D accumulation pattern, we reorganized the data as AO and BO groups to analyze the effect of loading concentration variations on the intensity and phasor FLIM events. Kruskal-Wallis ANOVA comparison of AO and BO organoids incubated with NP D, showed a statistical difference (at p level 0.05) of both intensity and FLIM signal events between groups with the similar tendency of median parameter value proportional to the concentration of nanoparticles (Tables S5-S8). Importantly, we did not detect phasor FLIM events in ‘no NP’ control organoids (40 organoids analyzed in total from both experimental replicates R1 and R2) with the chosen analysis settings, which made a rare event count value more significant (Fig. S10, Fig. 4C,D; Tables S3-4). In contrast, the intensity values of the ‘no NP’ control organoids (both AO and BO) overlapped with the intensity values of the low concentration groups (0.01 – 1 μg/mL; Fig. 4C, left panel). Based on the ‘reliability’ of one event detection we assumed that phasor FLIM can be used to count rare events of MNP accumulation in organoids. Thus, we counted the probability of organoid loading with NP D (a percentage of organoids with at least one detected phasor FLIM event from a total number of imaged organoids) as a function of NP D concentration in the surrounding media and determined that the probability of uptake per organoid increased with the concentration of NP D (Tables S3-S4). However, BO organoids were able to reach almost 100% loading probability at lower NP D concentrations (∼ 0.1 μg/mL) than AO organoids (∼ 10 μg/mL). This suggests the existence of intrinsic mechanisms ‘protecting’ the intestinal epithelium from NP D uptake from the apical membrane. At the same time, the heterogeneity in loading efficiency among AO organoids advocates the involvement of another factors, e.g. cell composition or metabolic state, facilitating NP uptake from the apical side. From a global perspective, the described event-based detection and analysis of heterogeneous NP uptake in the polarity-reverted enteroid culture can be viewed as an advanced approach with a potential to reflect a complex and perhaps competitive interactions of MNPs in the intestinal tissue.

### Phasor FLIM method enables analysis of complex mixtures of MNP in organoids

Real-world MNPs often display strong (poly)dispersity and a complex chemical composition, differing not only in the type of polymer backbone but also in their (partial) modifications and adsorbed ‘non-polymer’ components^47^. In addition, human and animal tissues can be exposed to multiple types of MNP at the same time. We therefore looked if our phasor FLIM event counting approach could discriminate between different nanoparticle types within the same organoids. Pristine dye-doped nanoparticles displayed a predictable and ‘calibrated’ fluorescence lifetime linear trajectory on their phasor plots (Fig. 1), while the phasor FLIM event counting reliably reflected the number of internalized nanoparticles (Fig. 4, S10). Thus, our goal was to utilize both properties to quantitatively trace a mixture of NP with distinct fluorescence lifetime values (NP D and NP B) in organoids.

First, we confirmed that internalized nanoparticles composed of pure NP species and their mixture could be clearly distinguished on a phasor plot. We compared the average G and S coordinates of phasor cluster points obtained from organoids loaded with 10 μg/mL NP D, NP B and their 1:1 mixture and detected a significant difference between their values, allowing to claim that these coordinates could be used as a marker of the internalized nanoparticles composition in organoids (Fig. 5A). Plotting of the representative cluster points with averaged coordinates demonstrated the existence of three lifetime zones on a phasor plot corresponding to pure D and B NP species and the intermediate ‘B + D’ zone, similarly NP in solution (Fig. S2). However, a real phasor cluster consists of many points each reflecting a NP uptake event and is potentially affected by the local NP environment and noise. Pure NP species cluster points deviate from the ‘center of a cluster’ within a distinct lifetime range (Fig. 5B), while mixed NP cluster points can be spread across all three lifetime zones, reflecting the proportion of the pure NP species presented simultaneously at the same pixel of the organoid image (Fig. 5D). To be able to count the percentage of intermediate cluster points in different lifetime zones of the phasor plot, we propose to apply a ‘lifetime fingerprint’ concept to determine range of the zones. Fig. 5 B-D demonstrates the step-by-step implementation of this approach for analysis of organoids loaded with the NP mixture. The fingerprint clusters of pure D and B species were reconstructed from a sum of corresponding individual clusters (7-10 per each group), allowing to increase the number of point units and providing better points diversity. For simplicity we chose to use only G coordinates as a representative of τ_ϕ_ values to set fingerprint zones for NP D and NP B. Accordingly, an average G ± standard deviation (SD) value was calculated as 0.4646 ± 0.0628 for NP D and 0.7056 ± 0.0524 for NP B and used as a threshold to determine the range of the intermediate zone (Fig. 5C). Thus, zone ranges were set as: G ≤ 0.5275 for NP D, G ≥ 0.6532 for NP B and 0.5275<G<0.6532 for the intermediate zone (B+D). The percentage of fingerprint cluster points detected in the intermediate zone did not exceed 15% from the total amount of the individual NP types, showing that most of the fingerprint points were in the range of a corresponding lifetime zones (Fig. 5C). Using G coordinate ranges we classified and calculated the percentage of points from ‘B+D’ clusters (Fig. S11, Table S9). Examples of individual organoid ‘fingerprint’ phasor plot clusters with point classification analysis are shown in Fig. 5D. The false color mask applied to the intensity image of internalized nanoparticles corresponds to the selected ROIs on the phasor plots and reflects the composition of the nanoparticles. Interestingly, organoids differed in their accumulation of NP D or B types, as evidenced by different cluster point percentages in the pure D and B lifetime fingerprint zones (Table S9). Some organoids (e.g. B_D_12_2, B_D_12_3 and B_D_16) demonstrated a high percentage of points (>50%) in the ‘B+D’ zone, related to accumulation of different NP types in the same vesicles or endosomes, probably due to utilization of the same internalization mechanism^48,49^. However, some organoids accumulated mainly one type of NP (e.g. organoids B_D_12_1, B_D_22 and B_D_14_2), as evidenced by cluster point percentages reaching up to 70-80% in some fingerprint zones. These results indicate the existence of at least 2-3 individual pathways of NP D and B internalization leading to accumulation of NPs in different vesicles and/ or reflects the difference in uptake related to cell composition of the organoids. We compared accumulation of the different NP types between AO and BO organoids and did not detect differences (Fig. S11) within each lifetime zone between AO and BO organoids, however, we observed a tendency among BO organoids to accumulate NP D (p=0.085, U = 50 vs. critical U value 45).

**Figure 5.**
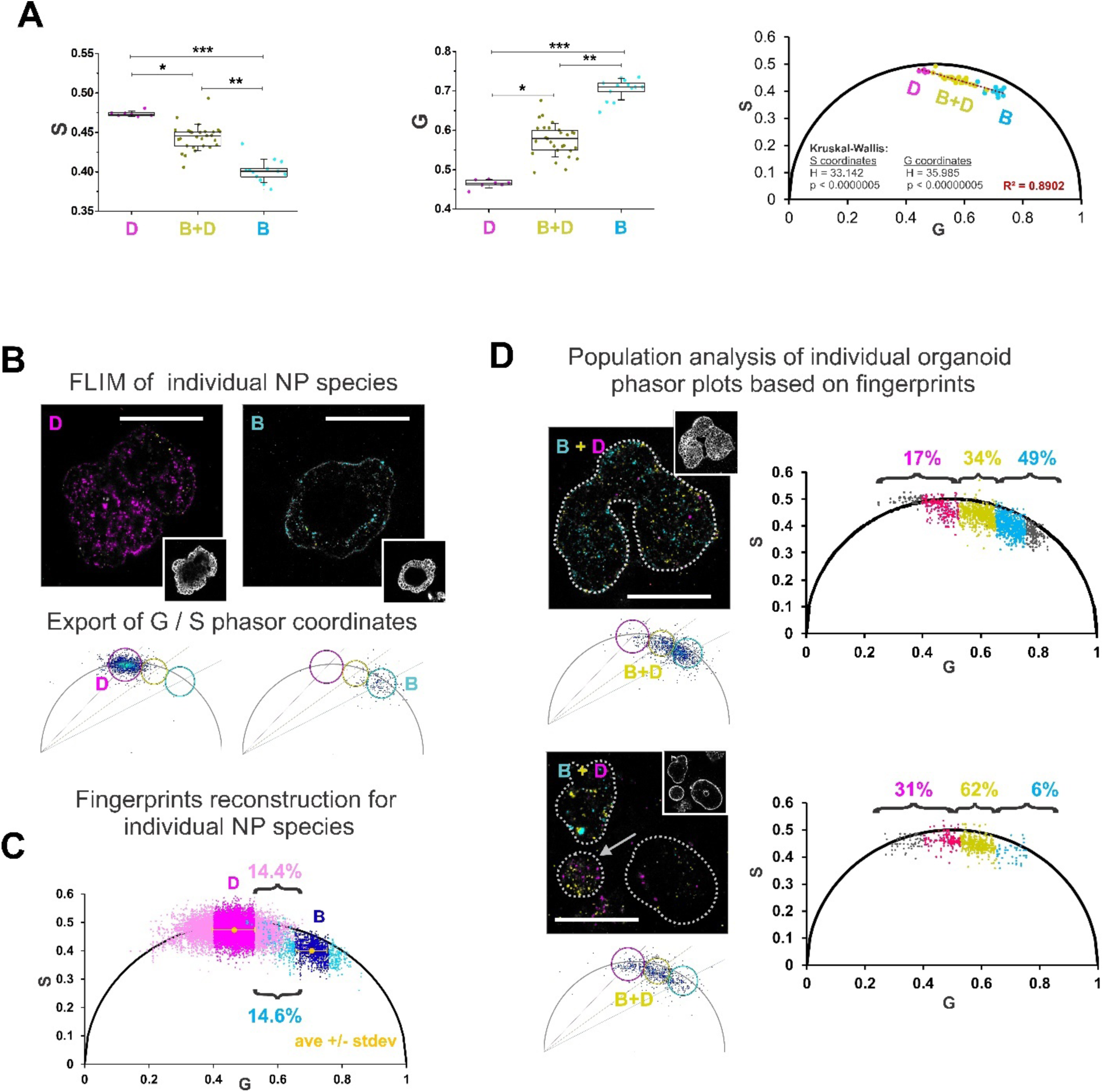
Phasor FLIM event counting approach resolves heterogeneous MNP populations in intestinal organoids. **A:** Distinct positions of NP D and NP B individually and their mixture in organoids on a phasor plot. From left to right: comparison of average S and G coordinates of phasor cluster groups, phasor plot. Box plots represent 25 and 75 percentiles, while whiskers show standard deviation. Each point corresponds to average G or S values of individual organoids. Asterisks indicate statistical difference between groups detected with the Dunn’s post-hoc test (*, p < 0.05; **, p < 0.0005; ***, p < 0.0000005). **B,C,D:** Illustration of the main steps of the fingerprint phasor FLIM event counting approach applied for tracking of heterogeneous MNP in organoids. **B:** Typical examples of NP D and NP B loaded BO organoids with corresponding phasor plots reconstructed in LAS X software. **C.** Phasor plot with fingerprint clusters of NP D (pink) and NP B (cyan) in organoids reconstructed in Excel from exported G and S coordinates. Yellow color indicates average cluster coordinates, magenta and dark blue color indicate the range of G_ave_ ± SD. **D:** Analysis of the NP uptake from the mixture of NP D and NP B in individual organoids with apical-basal topology. *Left:* phasor plots of individual organoids, reconstructed in Excel. The numbers show the percentage of total cluster points in NP D, D/B and B lifetime zones. Magenta and cyan color indicate cluster points in the range of corresponding G_ave_ +/- SD; yellow indicates D/B zone. All MNP intensity images have pseudocolor masks (magenta, yellow and cyan) based on the circular ROIs of the LAS X-produced phasor plots (NP D zone with τ_ϕ_ = 2.1 ns in magenta, intermediate D/B zone with τ_ϕ_ = 1.5 ns in yellow, and NP B zone with τ_ϕ_ = 1.0 ns in cyan, circle radius range is 46-36). Grayscale (bottom right) inserts show WGA labeling in the corresponding organoids. Scale bar is 100 µm.

In summary, we confirmed that the fingerprint cluster classification approach can be used for the analysis of intestinal organoids exposed to a complex mixture of MNPs, including the internalization pathways and possible synergism of nanoplastics during their uptake.

### Short-term exposure with nanoplastic particles does not affect cell energy production, mitochondrial function and inflammatory response

Lastly, we looked at how the potential physiological effects of the MNP could be assessed within our experimental settings. Within the intestinal epithelium and organoids, MNP can affect a multitude of processes, including the functions of the stem cell niche, absorptive and secretory cells, cell signaling and host-pathogen interactions^50^. We chose NP D (PS-MA) for these analyses, as they showed efficient accumulation in pig small intestinal organoids from the basal and apical sides.

Energy production is of paramount importance for any living cell. In eukaryotes, mitochondria play a vital role in this process. To determine whether NPs can affect mitochondrial homeostasis, we employed tetramethylrhodamine methyl ester (TMRM), a standard mitochondrial membrane potential probe^51^, to visualize mitochondrial polarization via FLIM^18,52^. When used in the intensity mode, TMRM only labels active mitochondria, however, this can be complemented by FLIM, visualizing lower mitochondrial polarization with longer lifetimes and more active mitochondria with shorter lifetimes^52^ (Fig. 6). We cultured organoids with and without NP D for 1 and 3 days, followed by WGA and TMRM labeling and FLIM. Intensity images confirmed WGA and TMRM labeling in the control organoids, and merged signal of WGA, TMRM and NP D in the NP treated organoids (video S1 and S2). Fast-FLIM images capturing TMRM lifetime distribution across all experimental conditions, with corresponding phasor plots for each group are presented in Fig. 6A. To look deeper in the effect of NP D on the fluorescence lifetimes of TMRM, we extracted pixel information (G, S coordinates) from phasor plots using the napari-FLIM-phasor-plotter (video S3), calculated mean G and S coordinates for each organoid and converted them into mean TMRM lifetime (Fig. 6B, right panel). While we found no significant differences in the TMRM lifetimes between the experimental groups over a 3 day period, we observed a downward trend in lifetime. While the reduced value in lifetime was small (NP D group: 2.075 ± 0.073 ns; control group: 2.236 ± 0.102 ns; Δ = 0.16 ns), it might indicate a potential increase in mitochondrial membrane polarization in organoids after 3 days of NP D treatment (Fig. S13A). It should be noted that different cell types in organoids are expected to display differences in cell cycle and mitochondrial activity^53–55^. It is therefore important to perform cell-specific segmentation for different cell types for such an analysis, to account for subtle changes of cell-specific metabolism upon MNP exposure.

**Figure 6.**
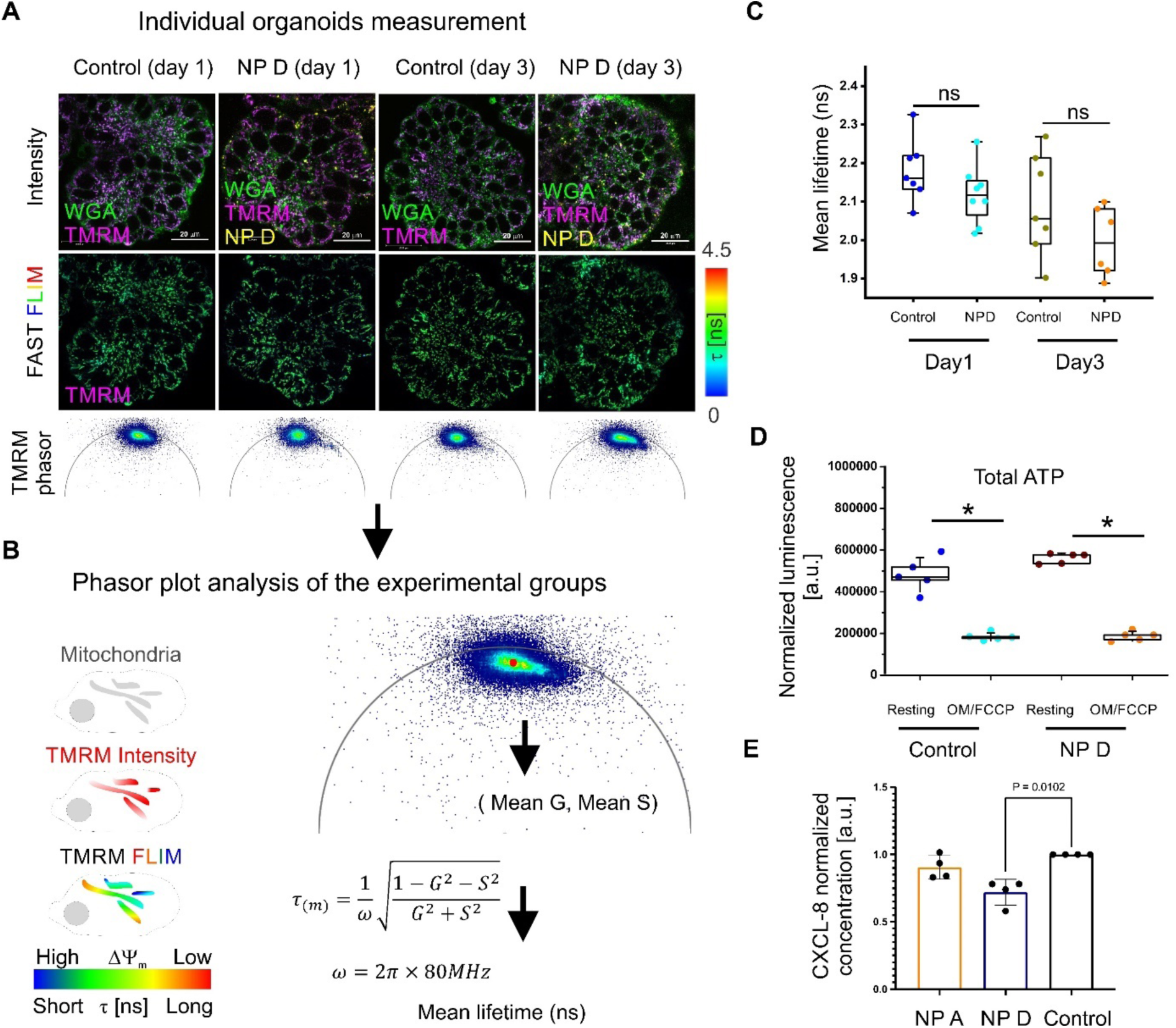
Assessment of physiological impact of absorbed NP D on mitochondrial polarization, cell energy budget and inflammatory cytokine expression. **A-C:** Workflow to investigate temporal effect of NP D exposure on the mitochondrial membrane potential (ΔΨ_m_) in BO organoid. **A:** Representative intensity images of basal-out organoid (BO) co-stained with WGA and TMRM, together with fast-FLIM images and phasor plots for TMRM, after 1- and 3-days NP D exposure. **B:** Scheme illustrating measurement of ΔΨ_m_ with FLIM of TMRM (left) and used quantification algorithm for the mean lifetime (right). C: Results of quantification of TMRM mean lifetime in control and NP D-treated BOs. **D:** Effect of NP D exposure on total cellular ATP in organoids. OM-FCCP indicates organoids pre-treated with 7 μM oligomycin, 1.4 μM FCCP. **E:** ELISA analysis of CXCL-8 expression in organoids after exposure to NP D (48 h). Data represent the mean ± SD of N=4 biological independent animals with 2 repeat experiments, analyzed with Kruskal-Wallis test. GP:0.1234(ns). 0.0332(*).

Since mitochondria are mobile organells^56,57^, we also looked at the applicability of the live mitochondrial tracking tool Nellie for our experimental setup (Fig. S12A). However, dynamic analysis of linear and angular velocity, length, and mitochondrial area did not show reproducible trends in three replicates (Fig. S12B-C). For instance, while the first replicate displayed reduced mitochondrial area in NP D-treated organoids compared to controls at day 3, the second replicate showed no difference and the third replicate showed increased mitochondrial area in NP D-treated organoids. Such inconsistencies can be a result of technical limitations of the used imaging approach (XYt scanning confocal microscopy with ‘fast FLIM’), not grasping mitochondrial dynamics in 3 geometrical dimensions (i.e. XYZt). Indeed, we observed rapid mitochondrial movement in our organoid model (video S4) with an estimated linear velocity of 0.02∼0.03 μm/s. At our optimized image acquisition settings (1024×1024 pixels, 400 Hz), each single-plane image acquisition required 2.59 s, while multi-sequential Z-stack of 5 depth coordinates would require 13 s for one image sequence, being too slow for tracking mitochondria in 4D. Potential future studies and development of live light-sheet FLIM-compatible systems would help addressing this issue, at least with the presented model^19,57–60^. Additionally, our mitochondrial tracking approach did not account for such aspects of functional complexity of apical-out organoids, as the cellular composition and the different cell cycle stages.

With no clear trend observed for the effect of NP D on mitochondrial polarization and mobility, we measured total ATP in NP D-treated and control organoids (Fig. 6D). While we forced the OxPhos to increase by using a low glucose medium^53,61,62^, we did not observe overall changes in ATP at resting and under mitochondrial uncoupling conditions, with oligomycin and FCCP treatment. Taken together, these results also confirm that the selected ‘pristine’ MNP at the chosen range of incubation conditions, does not affect overall mitochondrial mobility and cell energy budget in the majority of cells in the gut organoids. However, our analysis did not account for cell-type specific effects, which would be possible by adding live imaging tracers, such as markers of enterocytes, stem cells and others, and cell-specific segmentation^63^.

Intestinal epithelial cells not only provide a barrier function but also communicate with underlying immune cells. Even in a steady state, the gut epithelium secretes chemokines and cytokines to attract and inform immune cells. For instance, CXCL-8, a chemokine involved in neutrophil migration, is constitutively secreted by intestinal epithelial cells and is increased in response to pathogens or their virulence factors^64,65^. To determine whether exposure of gut organoids to NP influenced the communication network of the gut epithelium, we assessed CXCL-8 secretion levels in control and NP D and A treated organoids. As shown in Figure 6D, type D nanoparticles seemed to decrease CXCL-8 secretion levels by gut organoids. This can potentially result in reduced neutrophil numbers in the gut tissues and ultimately to an impaired ability to clear enteric infections.

Collectively, we illustrate here that the presented experimental model is compatible with functional assessment using live FLIM, dynamic intracellular organelle tracking and ‘bulk’ downstream assays, such as ELISA and ATP luminescent assays. This opens possibilities for a range of additional multi-parameter readouts, such as fluorescent cell sorting, single cell sequencing, mass spectrometry imaging, multiplexed immunofluorescence and others.

## Conclusion

Here, we developed a new approach, based on the use of bright NIR pristine nanoparticles, optimized dynamic imaging of live polarity-reverted small intestinal organoids and improved phasor FLIM-based inter-organoid tracing, interactions with different plasma membrane domains and quantification. This methodology enables deeper understanding of the structure-activity relationships of MNP matter interaction with tractable *in vitro* and potentially multi-organ-on-a-chip and *in vivo* live tissue models. The presented ‘lifetime barcoding’ feature allows for detecting different types of MNP and potentially their interactions with non-plastic matter, e.g. ions, biomolecules or metals. The potential limitation of presented approach can be in the need of designing ‘pristine’ nano- and microparticles for each tested type of MNP but in principle this can be also applied to real-world MNPs, pre-labeled with dyes, such as Nile Red. Importantly, the accurate choice of the labeling dye can benefit sensitivity, reliability and a ‘bar coding’ resolution of the proposed FLIM event counting approach, stemming from high quantum yields, NIR spectral properties and a well-characterized fluorescence lifetime response to environment. We have also illustrated application of methodology for assessing physiological effects on intestinal organoids, using dynamic live confocal and FLIM microscopies and downstream assays, such as ATP analysis and ELISA, meaning that it can be compatible with flow cytometry, -omics, multiple immunofluorescence and other downstream assays.

Remarkably, even with the limited selection of tested MNP, we found species-specific accumulation (different between mouse and pig) and minor effects on cytokine activation and mitochondrial polarization. This emphasizes the importance of single cell dynamic live microscopy imaging and specific cell type segmentation approaches, over the ‘bulk population’ assays, which can mask the cell-specific effects of MNP on the living cells.

## Materials and methods

### Polymers

Composition and properties of the co-polymers Eudragit RL100, PMMA-MA, PS-PVP and PS-MA were reported previously^26,30,66,67^.

### Small intestinal organoid culture materials

Lipidure^TM^-CM5206 (AMS.52000034GB1G, Amsbio, UK), Matrigel growth factor-reduced (734-0269, VWR, Belgium), human intestinal organoid growth medium (STEMCELL technologies, 06010, Belgium), mouse intestinal organoid growth medium (STEMCELL technologies, 06005, Belgium), DMEM high glucose GlutaMax^TM^ Supplement media (61965026, Gibco, Belgium), 0.5 M EDTA solution (15575020, Invitrogen, Belgium), 24-well Tissue Culture plates (734-2325, VWR, Belgium), sodium valproate (P4543, Sigma-Aldrich,Ireland), CHIR99021 (SML1046, Sigma-Aldrich,Ireland), PBS (18912-014, Gibco, Belgium).

Microscopy multi-well dishes were assembled using cover glass and silicon multi-well frame (cover glass with no. 1.5 thickness, e.g. μ-slide 12-well, Ibidi GmbH, Germany, or equivalent). Imaging was performed in phenol red-free DMEM (Sigma-Aldrich, D5030), supplemented with 10 mM glucose (G8270, Sigma-Aldrich), 1 mM pyruvate (11360-070, Gibco, Belgium), 2 mM GlutaMAX^TM^ (35050-038, Gibco, Belgium) and 10 mM HEPES-Na pH 7.2 buffer (15630-080, Gibco, Belgium) referred as imaging media (IM).

### Fluorescent probes

Alexa Fluor 488-conjugated Wheat Germ Agglutinin (WGA; W11261, Invitrogen, Belgium), Fluorescein-labeled *Ulex europaeus* Agglutinin I (UEA I; VEC.FL-1061-5, Vector Laboratories, Belgium), Nile Red (72485-100MG, Sigma-Aldrich, Belgium), MitoTracker Green (M7514, Invitrogen, Belgium), MitoTracker Red (M22425, Invitrogen, Belgium), Tetramethylrhodamine, methyl ester (TMRM) (T668, Invitrogen, Belgium), phalloidin-Alexa Fluor 546 (Invitrogen, A22283) Texas Red-X phalloidin (T7471, Invitrogen, Belgium).

### Production and characterization of pristine nanoplastics (NP)

Eudragit RL100, PMMA-MA, and PS-MA nanoparticles were obtained via nanoprecipitation according to previously published procedures^26,30,66,67^. Briefly, dibutoxy-aza-BODIPY dye^29^ and the polymer were dissolved in a water-mixable organic solvent, water was added rapidly under stirring and the organic solvent (along with most of water) was removed under vacuum. We used tetrahydrofuran:acetone (1:3 v/v) to dissolve PS-MA, tetrahydrofuran:acetone (1:9 v/v) for PMMA-MA (polymer first dissolved in tetrahydrofuran and the solution diluted with acetone) and acetone for RL100. The concentration of the polymers in the organic solvents was 0.004% wt. in all cases. We used 0.75% wt. of the dye in respect to the polymer for the staining. The dispersions after removing of organic solvents were concentrated to ∼1.5-2 mg /mL of the polymer content.

PS-PVP beads are commercially available and were stained via swelling in tetrahydrofuran:water mixtures as reported previously^30^. PS-PVP beads were doped with 0.75% wt. of the dye.

The size distribution and morphology of the nanoplastics were assessed using transmission electron microscopy (TEM) as described previously^27^. Briefly, NP were drop cast (2 µl, 20 µg/ml) and dried overnight at room temperature (RT) on Formvar/Carbon-coated hexagonal copper mesh grids (FCF200H-CU-TB, Electron Microscopy Sciences). Nanoplastics (n=150) were observed on a transmission electron microscope JEM 1010 (Jeol, Ltd, Japan) equipped with a charge-coupled device side-mounted Veleta camera (Emsis, Germany). Nanoparticle size was measured manually using ImageJ software (NIH, USA) and graphs were made using Graph Pad Prism 9.

The fluorescence excitation and emission spectra were measured on a Fluorolog-3 luminescence spectrometer (Horiba, Germany) equipped with a NIR-sensitive R2658 photomultiplier (Hamamatsu, Germany) in a quartz cuvette as described previously^27^. The fluorescence lifetime of NP in water, fetal bovine serum (FBS) and other solutions were measured on a Stellaris 8 Falcon FLIM microscope (see section ‘Microscopy’ below).

### Pig and mouse small intestinal organoid cultures

Pig small intestinal organoids were produced from the jejunum (Table S10) as described previously^64^. Organoid lines used in the experiments were developed from 3 individual animals. Three organoid lines (porgj2, porgj3, porgj4) were used for studying MNP interactions with the intestinal epithelium. The lines porgj2, porgj3, pigD, pig13 and pig24 were used for assessing effects on chemokine expression. The porgj2 line was used for total ATP, mitochondrial polarization and mitochondria mobility analysis. Validation of phasor plot based counting method and MNP co-loading analysis were done on porgj3 line. Mouse small intestinal Lgr5-GFP organoid line (Lgr5-EGFP-ires-CreERT2)^43^ was provided by J. Puschhof and H. Clevers (Hubrecht Institute, Utrecht University, The Netherlands).

The mouse and pig organoids were cultured at density 150-300 organoids per 50 μL Matrigel dome in 500 μL of organoid growth medium per well of 24-well plate. Human IntestiCult organoid growth medium HGM (STEMCELL technologies, 06010, Belgium) was used for culturing pig organoids^64,68^. Mouse IntestiCult organoid growth medium MGM (STEMCELL technologies, 06005, Belgium) supplemented in house with 1 mM sodium valproate and 3 μM CHIR99021 was used for Lgr5-GFP mouse intestinal organoids^53^. To prevent accumulation of dead cells in the organoid lumen and growing large structures, organoids were passaged by mechanical disruption of Matrigel domes, as described previously^69^. The disrupted organoid suspension was diluted to 10 mL with washing medium (DMEM high glucose GlutaMax^TM^ Supplement, Gibco, 61965026, Belgium), collected by centrifugation (300g, 5 min, 4°C), the supernatant was gently aspirated and the pellet was resuspended in an appropriate volume of liquid ice-cold Matrigel for passaging at a 1:3 ratio. The organoid/Matrigel mixture was dispensed (50 μL/well) in a pre-warmed 24-well plate, incubated for 5 min at 37°C to solidify Matrigel domes and, subsequently, growth medium (500 μL/well) was added to cover the Matrigel domes. Typically, organoids were passaged every 3-4 days.

### Polarity reversion of organoids

#### Preparation of low-attachment plasticware

Low-attachment plasticware is essential for polarity-reverted ‘apical-out’ (AO) organoids: (1) to prevent adherence of the BO organoids to the plastic culture plates and loss of 3D morphology; (2) the prevent the non-specific adhesion of organoids to the plastic surfaces during handling. To prepare low-attachment centrifuge vials and 24-well culture plates, 0.5% lipidure-coating solution (95% molecular grade ethanol, made from the powder Amsbio, AMS.52000034GB1G) was added to cover the surface, followed by 1-5 min incubation at RT, under sterile conditions. Subsequently the coating solution was aspirated and the coated plastic surfaces were dried under a sterile air, and stored at RT until use.

#### ‘Apical-out’ (AO) organoid suspension culture in low-attachment plates

To produce AO organoids, we adapted the protocol of Co et al^38^. Briefly, 1-3 day old cultures of BO organoids in 50 μL Matrigel domes at density of ∼150-300 organoids/dome were rinsed once with ice-cold 5 mM EDTA-PBS solution, followed by the dislodging of the Matrigel domes and resuspension by pipetting with 1 mL pipette tips (low binding tips; WB 5174S, Westburg) in a ice-cold 5 mM EDTA-PBS solution (500 μL/dome). Subsequently, the Matrigel/organoids suspension (pooled from 4 domes) was transferred into lipidure-coated 15 mL centrifuge vials and the volume was adjusted with cold EDTA-PBS solution to 12 mL. Thz suspension was incubated for 1 h at 4°C with continuous gentle mixing to completely dissolve the Matrigel. Organoids were collected by centrifugation (300g, 3 min, 4°C), washed once with 10 mL ice-cold washing medium (DMEM high glucose GlutaMax^TM^ Supplement), followed by resuspension of the pellet in 3.2 mL of 37°C pre-warmed HGM and dispensed in lipidure-coated (low-attachment) 24-well plate (400 μL per well). Organoids were cultured for 18-20 h to induce polarity reversion in the absence of basement membrane signaling. AO organoid suspension culture could be cultured for up to 5 days.

#### Loading of AO organoids with Dibutoxy-aza-BODIPY-doped nanoparticles and staining with fluorescent dyes

To evaluate the uptake of MNPs (Table 1), nanoparticles A-D were added to 1 day old AO organoid suspension culture in a low-attachment 24-well plate in HGM (AO organoid density at ∼ 100-150 organoids/well, 400 μL medium, 10 μg/mL MNP) and incubated for 18-24 h prior to microscopy analysis.

For analysis of mitochondrial mobility, polarization, chemokine secretion and ATP, NP D was used at 10 μg/mL. To improve the yield of loaded organoids in these experiments the loading was performed simultaneously with the polarity reversion. For the phasor-based counting method validation, NP D were added to organoids during the reversion procedure at 0, 0.01, 0.1, 1, 10, 25, 50 and 100 μg/mL in medium. For co-loading experiment, NP B and D were used individually (for a fingerprint analysis) and together at 10 μg/mL with the loading procedure done during the polarity reversion.

For the time lapse loading experiments, NP D (10 μg/mL) were added directly during monitoring to AO organoid suspension culture in “imaging medium” or IM (DMEM D5030 Sigma-Aldrich, without phenol red and sodium bicarbonate, supplemented with 10 mM D-glucose, 2 mM GlutaMax, 1 mM sodium pyruvate, 10 mM HEPES-Na, pH 7.2). Microscopy experiments were performed at different time points (0.5-24 h).

Where indicated, a 1 h co-staining with different fluorescent probes (see Table 2) was performed in HGM. Prior to the microscopy, organoids were washed 3 times to remove nanoparticles and other fluorescent probes by media exchange with the IM. For TMRM-FLIM^52^, 10 nM of TMRM was added to IM after the final media exchange and remained in solution during the imaging procedure.

#### Organoids fixation with PFA and fluorescent phalloidin labeling of F-actin

For phalloidin-based F-actin labeling of apical membranes, intestinal organoids (AO organoid suspension or BO organoids embedded in Matrigel) were washed 3 times with pre-warmed (37°C) PBS, immediately fixed with warm 4% paraformaldehyde in PBS (10 min, RT), washed 5 times with PBS, permeabilized with 0.1% Triton-X100 in PBS (10 min, RT), incubated with blocking solution (5% FBS in PBS) and subsequently stained with Alexa Fluor 546 or Texas Red-conjugated phalloidin-conjugates in blocking solution (1 h, RT). Stained organoids were washed 5 times with PBS and stored in PBS at 4°C until analysis. The organoids were not embedded between glass slides to preserve their 3D organization.

For NP D concentration-dependent uptake study, organoids were fixed with 4% PFA as described above and stored at 4°C in PBS until the analysis by microscopy.

#### Microscopy

Confocal FLIM microscopy was performed on a Stellaris 8 Falcon microscope, (Leica Microsystems, Ghent Light Microscopy Core, Ghent University), equipped with a white-light laser (440−790 nm), HC PL Apo 10×/0.4 air, HC Fluotar 25×/0.95 W, HC PL Apo 40×/1.25 GLYC corr, HC PL Apo 63×/1.4 oil objectives, HyD X, HyD R and HyD S detectors, temperature- / CO_2_-controlled incubator, and dedicated LAS X acquisition and analysis software (ver. 4.6.0), as described previously^27,72^. For routine microscopy, a 40×/1.25 GLYC corr. objective was used, with typical settings: 80 MHz white-light laser frequency pulse, scan speed 100-400 Hz, pixel dwell time 2.087 – 7.688 µs, pinhole 1.0 – 1.2 airy units, 1-3 frame repetition rate, 512 x 512 or 1024 x 1024 pixels resolution. The following general excitation and emission settings were used for fluorescence imaging: dibutoxy-aza-BODIPY-doped nanoparticles (MNP) excited at 698 nm (laser power 10%, emission 717 - 753 nm), TMRM excited at 540 nm (laser power 1%, emission 559 - 585 nm), WGA-Alexa Fluor 488 excited at 490 nm (laser power 4%, emission at 495 - 520 nm), UAE-Fluorescein conjugate excited at 495 nm (laser power 10.54%, emission at 497-568 nm), Nile Red excited at 490 nm (laser power 4%, emission collected at 588 - 620 nm), MTR excited at 581 nm (laser power 1.2 %, emission collected at 594 - 654 nm), MTG excited at 490 nm (laser power 6.6 %, emission at 497 −551 nm), phalloidin-Texas Red conjugate excited at 579 nm (laser power 3.21%, with emission collected at 588 - 649 nm).

Widefield fluorescence microscopy was performed on an inverted fluorescence microscope IX81 (Olympus), equipped with motorized Z-axis control, CoolLED pE4000 (16 channels, 365-770 nm), ORCA-Flash4.0LT+ (Hamamatsu) cMOS camera, temperature controlled stage (OkoLab), and air objectives 4x/0.13 UPlanFLN, 10x/0.3 UPlanFLN, 40x/0.6 LUCPlanFLN, water immersion objective 60x/1.0 LUMPLFLN, CellSens Dimension v.3 software as described previously^73,74^.

#### FLIM analysis of dibutoxy-aza-BODIPY-doped nanoparticles in solution

Dispersions of different nanoparticle types (A, B, C and D) were prepared by diluting stock nanoparticle dispersions to a final concentration of 500 μg/mL in 25% FBS and 50% imaging medium. For phasor plot comparison in response to nanoparticle dilution, series of nanoparticle type D dilution in milli-Q grade water were performed over concentration range of 0.58-500 μg/mL. milli-Q grade water was used as a negative control. Subsequently, confocal FLIM measurements of a 30 μL drop of nanoparticles dispersion on a cover glass surface were performed with HC PL Apo 10×/0.4 air, pinhole 1.0 AU. and 1.16 x 1.16 mm field of view was imaged with 512 x 512 pixels resolution, 200 Hz scanning speed, 3 frame repetition per image. The frequency of the laser pulse was set to 80 MHz with the power of the laser excitation at 698 nm 8%. Pixel by pixel acquired fluorescence decays were automatically converted into phasor plot data in dedicated LAS X software (Leica Microsystems, version 4.6.0). The corresponding τ_ϕ_ (tau phase) of the phasor patterns was calculated from the center position of the circular region of interest (ROI) applied to the phasor plots. ROI-based color coding was applied to related pixels on the images. For comparing lifetimes of different nanoparticle types, the following phasor plot settings were used: first harmonic, threshold 7, median filter set as 3, bin 1, circular ROI radius 14. For the analysis of a dilution effect on nanoparticle type D lifetimes, the following phasor plot settings were used: first harmonic, threshold 4, median filter set as 11, nanoparticle signal circular ROI radius 58, with pixel binning 1 or 3.

For precise calculation of nanoparticle fluorescence lifetime, a 3-exponential reconvolution fitting model was applied to collected global decays and intensity weighted mean fluorescence lifetime (τ_m_) in LAS X 4.6.0 software (Leica Microsystems).

#### Comparison of intensity- and lifetime (phasor)-based analysis approaches

The experiment was repeated twice using the same organoid line. Organoids were loaded with NP D at various concentrations (0-100 µg/mL) during the polairty reversion and co-stained with WGA-Alexa Fluor 488 for determining topology and segmentation. Subsequently, they were fixed with 4% PFA, washed with PBS and stored at 4°C in PBS, prior to microscopy. FLIM microscopy was performed using HC PL Apo 40×/1.25 GLYC corr objective, 1024 x 1024 pixel resolution, 1 frame repetition, 100 Hz scanning speed, pixel dwell time 7.7 μs, excitation at 490 nm (1.8-3.3% laser power, for WGA) and 698 nm (10% or 30% laser power intensity, as indicated, NP D), 80 MHz frequency laser pulse, zoom 1, one sequence excitation and emission light collection with HyD X (range 495-541 nm) and HyD R (717-753 nm) detectors on a confocal FLIM microscope. Each experimental group including control (unloaded organoids) contained 17-33 organoid FLIM images, acquired as described above. First, FLIM images of NP D fluorescence and corresponding control images were pre-analyzed in LAS X software (Leica Microsystems, version 4.6.0) to confirm that only phasor clusters in the 1.96 ns lifetime zone were present and no unrelated phasor clusters appeared on the control phasor plots images. Different median filter and pixel binning values were tested to decrease dispersion of the phasor clusters with reasonable preservation of the image spatial resolution. After choosing pixel binning 3 as the setting parameter for the phasor plots we optimized the intensity (not phasor) threshold in randomly screened images from the experimental groups at the lowest and highest loading concentrations. The threshold was selected to filter out the blurring pixels from the images of the high NP D loading concentration groups, while keeping the main pixels, reflecting the real NP signals inside cells, intact. Pre-analyzed raw organoid images were exported in .PTU format and analyzed in napari viewer, with a FLIM phasor plotter plugin [<u>GitHub - zoccoler/napari-flim-phasor-plotter</u>], (<u>10.5281/zenodo.12620955</u>). Using WGA as a marker of organoid ROI selection the ROI mask was manually applied to the NP D fluorescence channel image and the ROI image was processed with application of pixel binning 3 and intensity threshold value 188 a.u. with the subsequent phasor plot reconstruction using median filter 10. Accordingly, all individual organoids on images were analyzed and their ROI area and total phasor plot points G and S coordinates were exported as .csv table format files. To speed up the analysis a macros code ‘Supplementary Code area_GS_export’ (<u>10.5281/zenodo.14742564</u>) was produced in Python using Claude AI (www.claudeai.com) and validated with manually extracted and processed data. The content of all .csv phasor plot coordinate files was pre-screened to remove nonsense point data (points where both G and S coordinates or one of them had value 0) and the total number of points from NP D lifetime zone was counted using Claude AI (‘Supplementary code_GScount’, <u>10.5281/zenodo.14742564</u>). Subsequently, all the exported data (apical-basal organoid topology, organoid ROI square area, number of cluster points) were organized in Microsoft Excel and the number of points was normalized per organoid ROI square area to have a number of FLIM events per square unit. Accordingly, organoids were classified based on their apical-basal topology into AO and BO (include BO and AB organoids) organoids groups for all loading concentration groups.

In addition to the FLIM and data analysis, the same organoid suspension sample was imaged with widefield fluorescence microscope with air UPlanFLN, 10x/0.3 and UPlanFLN, 40x/0.6 LUCPlanFLN objectives (Olympus-Evident) as described above. The following acquisition settings were applied: 1024×1024 pixel resolution, NP D excited with 660 nm (LED power 40%) with emission collected at 705-845 nm (exposure time 40 ms) and WGA-Alexa Fluor 488 excited with 470 nm (LED power 40%) with emission collected at 510-550 nm (exposure time 7.363 ms). Images were exported as .TIFF files and analyzed in Fiji software. The organoid ROI mask was made based on WGA-Alexa Fluor 488 segmentation and applied to NP D spectral channel fluorescence intensity image to extract total organoid intensity. The NP D background image intensity was calculated as an average from 10 random 1-pixel points surrounding organoids. Organoid ROI square areas, background and total organoid intensities were exported to Microsoft Excel table file and an individual organoid intensity values per area were calculated by substracting background intensity from the total organoid intensity value and normalizing by the ROI area.

Both FLIM events and intensity datasets were tested for normal distribution using normality test calculator [Georgiev G.Z., *"Normality Calculator"*, [online] Available at: https://www.gigacalculator.com/calculators/normality-test-calculator.php URL [Accessed Date: 24 Jan, 2025] using a sum of normality tests, including Shapiro-Wilk test / Shapiro-Francia test (n < 50 / n > 50), Anderson-Darling test, Jarque & Bera test, Cramer-von Mises test, d’Agostino-Pearson. The statistical analysis was done in OriginPro software with graphs made in Origin 6.0 (OriginLab). Accordingly, non-parametric tests were used: (i) Mann-Whitney for comparison between AO and BO organoids for each loading concentration data set and (ii) Kruskal-Wallis ANOVA for multiple group comparison with Dunn’s and Conover’s post-hoc test analysis between individual concentration groups among AO and BO grouped organoids.

#### Phasor-FLIM-event counting approach for analysis of MNP mixtures

FLIM imaging data of control (no NP) and organoids loaded with NP D and B nanoparticles (as a mixture or separately) were exported as .PTU files, processed with the napari-phasor-plotter plugin to extract G and S phasor point coordinates. After removing of nonsense points as described above, phasor coordinates were organized in Microsoft Excel table. The average values of G and S coordinates for individual NP D, NP B and MNP mixture-loaded organoids were counted and compared using Kruskal-Wallis ANOVA with Dunn’s post-hoc test. The fingerprint phasor clusters of NP D and NP B in organoids were reconstructed in Microsoft Excel from the sum of all corresponding phasor plots coordinates. For every fingerprint cluster the values of average G coordinates (G_fave_) ± standard deviation (SD) were calculated and applied as thresholds for the pure NP D and B fluorescence lifetime zones. Similarly, a few phasor clusters of individual organoids loaded with a MNP mixture were reconstructed and the percentage of pure D, pure B and mixed D/B intracellular MNP were counted by the attributing corresponding G coordinates to the following zones: G ≤ G_fDave_ + SD_D_ (NP D lifetime zone), G ≥ G_fBave_ – SD_B_ (NP B lifetime zone) and G_fBave_ – SD_B_ > G < G_fDave_ + SD_D_ (D / B mixed lifetime zone). The corresponding Claude AI generated code was used (‘Supplementary code_tau_zones’, <u>10.5281/zenodo.14742564</u>).

#### CXCL8 ELISA

CXCL8 was quantified in the culture supernatant of apical-out enteroids using a swine-specific CXCL8 DuoSet ELISA kit (R&D systems, Minneapolis, MN, USA) following the manufacturer’s instructions. Culture SN was diluted ½ in reagent diluent. Absorbance values (450 nm) were measured using a Tecan Spark and converted to concentrations using Deltasoft software. The CXCL8 concentrations were then normalized to the protein concentration of apical-out enteroid lysates for each condition. Protein concentrations were determined using BCA.

#### Total ATP measurements in organoids

This was performed using a CellTiter-Glo viability assay (Promega, G7591) with following modifications. Two independent experimental repeats were done on the same organoid line at different passages. Four domes of BO porgj-2 organoids were used to produce 4100 μL of AO organoid suspension in HGM. The suspension was dispensed at 200 μL per well in 20 wells of a flat bottom 96 well plate (Greiner) pre-treated with Lipidure^TM^ coating solution. Nanoparticles type D were immediately added to 10 wells in a final concentration 10 μg/ mL, leaving untreated 10 control wells and incubated for 48 h, followed by replacement of HGM with imaging media (IM) without glucose with sequential (6 times) partial medium exchange, resulting in final 200 μL of medium per well (this procedure allowed to decrease glucose in the media to activate OxPhos in organoids). Subsequently, organoids were incubated for 24 h prior to ATP measurement on day 4. Nanoparticle loading efficiency (the percentage of nanoparticle-positive organoids from total number of organoids analyzed, with partially loaded organoids counted as one loading event) was assessed on a widefield inverted fluorescence microscope IX81, 40x/0.6 LUCPlanFLN objective (Olympus). For total ATP measurement in organoids at rest, first 130 μL of organoid suspension were treated with mock (DMSO) or 1.4 μM FCCP/ 7 μM oligomycin mixture for (20 min, 37 °C, 5% CO_2_). Collectively, four different groups with 5 wells per group were tested: control / resting, control / stress, nanoparticles D treated / resting, nanoparticles D treated / stress. After treatment, suspensions from individual wells (140 μL) were collected into individual 500 μL lipidure-coated vials. To ensure complete collection of organoids, wells were additionally rinsed with 150 μL of no glucose IM, which was combined in the corresponding vials with the main organoid suspension. Organoids were collected by centrifugation (5 min, 300g, RT), and pellet was gently resuspended in 10 μL of media. Subsequently, 100 μL of ATP reagent (1:1 mixture of no glucose IM Cell Titer-Glo reagent) was added to individual vial and vigorously mixed (vortex) for 18 s. Vials were centrifuged (5 min, 300g, RT) and 80 μL of debris-free supernatant was collected into white opaque-walled 96 well plate (Corning), proceeded by the measurement of luminescence with Varioscan microplate-reader (Thermo Fisher Scientific) with 1000 ms exposure time. The background noise signal was determined as an average signal measured from 5 individual wells with 80 μL of media taken from the mixture of 10 μL of no glucose IM with ATP reagent. This was subtracted from a total luminescence signal of each individual experimental well and normalized per total average area square of organoid per treatment condition (measured by transmission light microscopy of individual sample wells of 96 well plate with widefield inverted microscope Olympus IX81).

Statistical analysis was done in OriginPro 12 (OriginLab) using Mann-Whitney non-parametric test to compare between groups of no NP (control) and NP D (loaded organoids with confirmed uptake) at resting (no stimulation) and FCCP / Oligomycin stimulation conditions. The n for each group was 5 wells with organoid suspensions.

#### Phasor plot analysis of MNP effects on mitochondrial polarization

To study the effect of NP influence on mitochondrial polarization in intestinal organoids, we exported TMRM FLIM data in PTU format from LAS X software and processed them using the napari-flim-phasor-plotter plugin (https://zenodo.org/records/12620956) and ClaudAI designed macros code ‘Supplementary Code 1’ (https://github.com/HangZhouFLIM/FLIM_nano). For each PTU file from control (no NP) and NP D groups (days 1 and 3), we selected the TMRM channel and manually cropped the organoid region to remove artifacts. Phasor analysis was performed (bin size=1, threshold=10, median filter=10) at 80 MHz laser frequency. G-S coordinates and ROI measurements were exported as .csv files. We then calculated the mean G and S from G and S coordinates, with data filtering (2.5-97.5 percentile outlier removal of G and S respectively) to remove extreme values, and converted to mean *τ*_(*m*)_ using the equation: *τ*_(*m*) = 1/*ω* √((1 − *G*^2 − *S*^2)/(*G*^2 + *S*^2)) with ω = 80 MHz and draw the boxplot in control (no NP) and NP D groups (days 1 and 3) using ClaudAI designed ‘Supplementary Code 2’ (https://github.com/HangZhouFLIM/FLIM_nano).

#### Mitochondrial dynamics and morphology analysis

To investigate impact on the mitochondrial dynamics and morphology, AO organoids (Porgj-2) were cultured for 24 h before NPD treatment, followed by additional 1- or 3-day cultures. At each timepoint, organoids were stained with WGA (1 h), washed three times with imaging medium, and then treated with TMRM for mitochondrial visualization. Live microscopy was performed using a Stellaris Falcon 8 confocal microscope (1024 × 1024 pixels, 400 Hz scan speed, excitation: 510 nm, emission: 555-650 nm) with 2-minute time-lapse acquisition yielding 47 sequential timeframes (t1 to t47). Images were exported as OME-TIFF format using LAS X software and preprocessed to separate WGA, TMRM, and NPD channels using custom Python scripts in napari (‘Supplementary Code 3’, https://github.com/HangZhouFLIM/FLIM_nano). Mitochondrial features (TMRM images) were tested using the Nellie plugin^75^ in napari (https://zenodo.org/records/13863809) for single timeframes, and multiple timeframes were processed through custom batch-processing scripts based on Nellie code from GitHub (‘Supplementary Code 4’, https://github.com/HangZhouFLIM/FLIM_nano) and exported as CSV files. Ten key parameters were selected: linear velocity, angular velocity, tortuosity, aspect ratio, length, solidity, extent, minor axis length, major axis length, and area, and data analysis was performed at organoid level (mean values of all mitochondria per organoid, visualized with box plots) comparing NP and no NP groups on day 1 and day 3 using custom analysis scripts (‘Supplementary Code 5’, https://github.com/HangZhouFLIM/FLIM_nano).

#### Data assessment and Statistics

Origin software (ver 6.0 and 12) was used for statistics, except for mitochondrial dynamics and morphology. For two group comparisons, Mann Whitney U test was used. Error bars and p values were reported in the figures and figure legends. Statistical analysis of mitochondrial dynamics and morphology was calculated using Python. We collected and analyzed all data objectively using instruments without bias and experimental samples were randomly chosen into groups.

#### Data and code availability

Raw PTU and CSV files for analysis of organoids loaded with MNP mixtures by Phasor-FLIM-event counting approach are available at Zenodo (<u>10.5281/zenodo.14742564</u>). Custom codes used for GS export and GS coordinates analysis are available at Zenodo (). Raw PTU files and processed CSV files for mitochondrial polarization phasor analysis in organoids are available at Zenodo (https://zenodo.org/records/14417182). Mitochondrial microscopy data, Nellie-processed csv files, and associated analysis plots generated using custom code are available at Zenodo (https://zenodo.org/records/14417688) and a public GitHub repository (https://github.com/HangZhouFLIM/FLIM_nano).

## Supporting information

Supplementary Figures and Tables combined

## Acknowledgments

This research was supported by the Special Research Fund (BOF) grants (BOF/STA/202009/003, BOF/BAF/1y/25/1/004), Research Foundation Flanders (FWO, I001922N, I004124N), and the European Union, fliMAGIN3D-DN Horizon Europe-MSCA-DN No. 101073507 grants.

We would like to thank Lars Vereecke (advice on lectins), Johannes Swinnen, Nina Ravoet, Max Nobis and Colinda Scheele for support with two-photon microscopy experiments, as well as Alain Labro, Chris L. Langsdorf and Winter Vandenberghe for sharing fluorescent dyes.

