## Supplementary Figures and Tables combined for "Fluorescence Lifetime Imaging Microscopy (FLIM) visualizes internalization and biological impact of nanoplastics in live intestinal organoids"

### Supplementary Information

#### Supplementary figures S1-S13

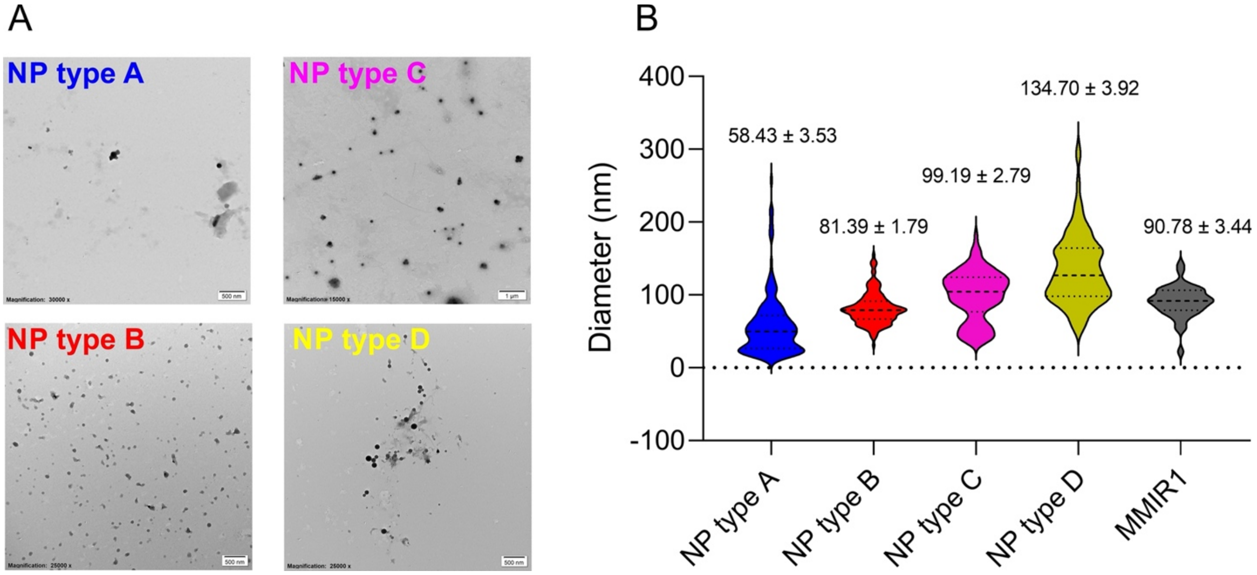


**Figure S1. TEM and size distribution of the NP A-D. A.** Transmission electron microscopy (TEM) of nanoplastic particles. Scale bar is 500 nm for type A, B and D and 1 µm for NP type C (1 µm). **B**. Size distribution of the nanoplastics type A-D with RL-100 based nanoparticles MMIR1 as control. Data shows the average size ± SEM for 150 counted nanoparticles and 50 for nanoparticle control.

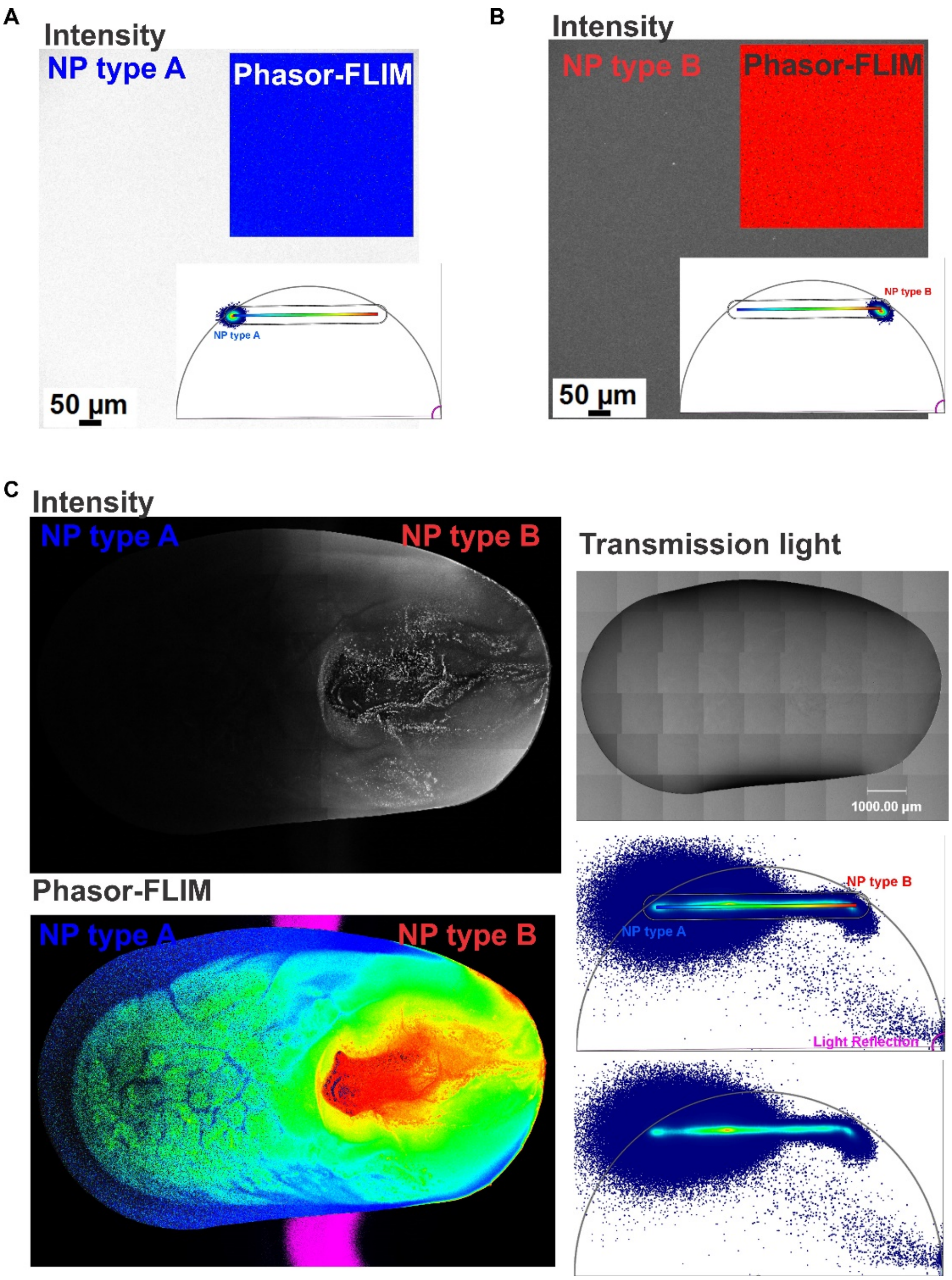


**Figure S2. FLIM resolves different types of NP in aqueous dispersions**. NP A and B were measured individually and in half-mixed water drops at 0.5 mg/ mL with the following analysis of the phasor patterns changes. **A, B:** FLIM of pure NP A and NP B species prior to mixing. Raw intensity images (at the same scale range) are shown together with the color mask applied intensity images. False color coding applied as shown on the corresponding phasor plots with a range of $\tau_\phi$ from 3.7 (blue) to 1 (red) ns. **C:** FLIM mosaic imaging of the half-mixed solutions of NP A and B. The raw intensity and phasor-mask applied intensity images are shown together with the corresponding transmission light image of the combined (half-mixed drops of NP A and B water solutions). Resulting mosaic phasor pattern (with the corresponding ROI mask – on the top and the raw phasor – on the bottom) localized on a line between opposite phasor positions of the individual unmixed NP A and B species.

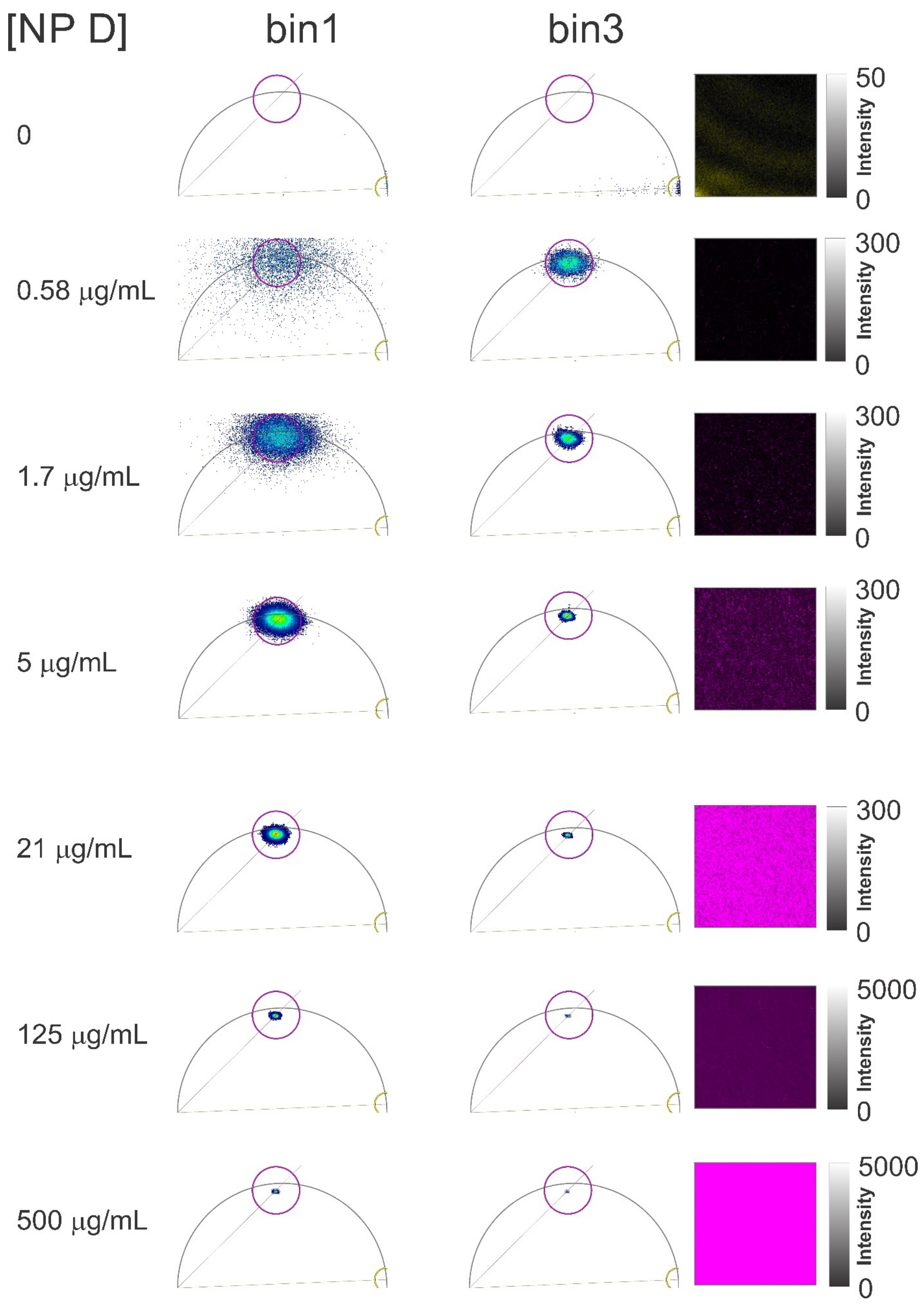

[NP D]
bin1
bin3
0
50
Intensity
0
0.58 μg/mL
300
Intensity
0
1.7 μg/mL
300
Intensity
0
5 μg/mL
300
Intensity
0
21 μg/mL
300
Intensity
0
125 μg/mL
5000
Intensity
0
500 μg/mL
5000
Intensity
0

**Figure S3. Pixel binning decreases scattering of the noise and NP D signal phasor patterns on a phasor plot.** The imaging was done with 200 Hz scanning speed, 3 frame repetition, 512x512 resolution, pinhole 1 AU. Phasor analysis of series of NP D dilutions in water was performed with LAS X version 4.6.0 (Leica Microsystems) software: harmonic 1, median filter 11, threshold 4, pixel binning 1 or 3. Magenta circular ROI marks the estimated localization of NP D related pixels ($\tau_\phi = 2.023$ ns, radius 58), yellow circular ROI marks the estimated localization of noise pixels ($\tau_\phi = 0.083$ ns, radius 28). Examples of fluorescence intensity images (for pixel binning 3) of noise (the water drop) and NP D dilutions presented on the right with the applied false color masks based on corresponding phasor plot ROIs (on the left).

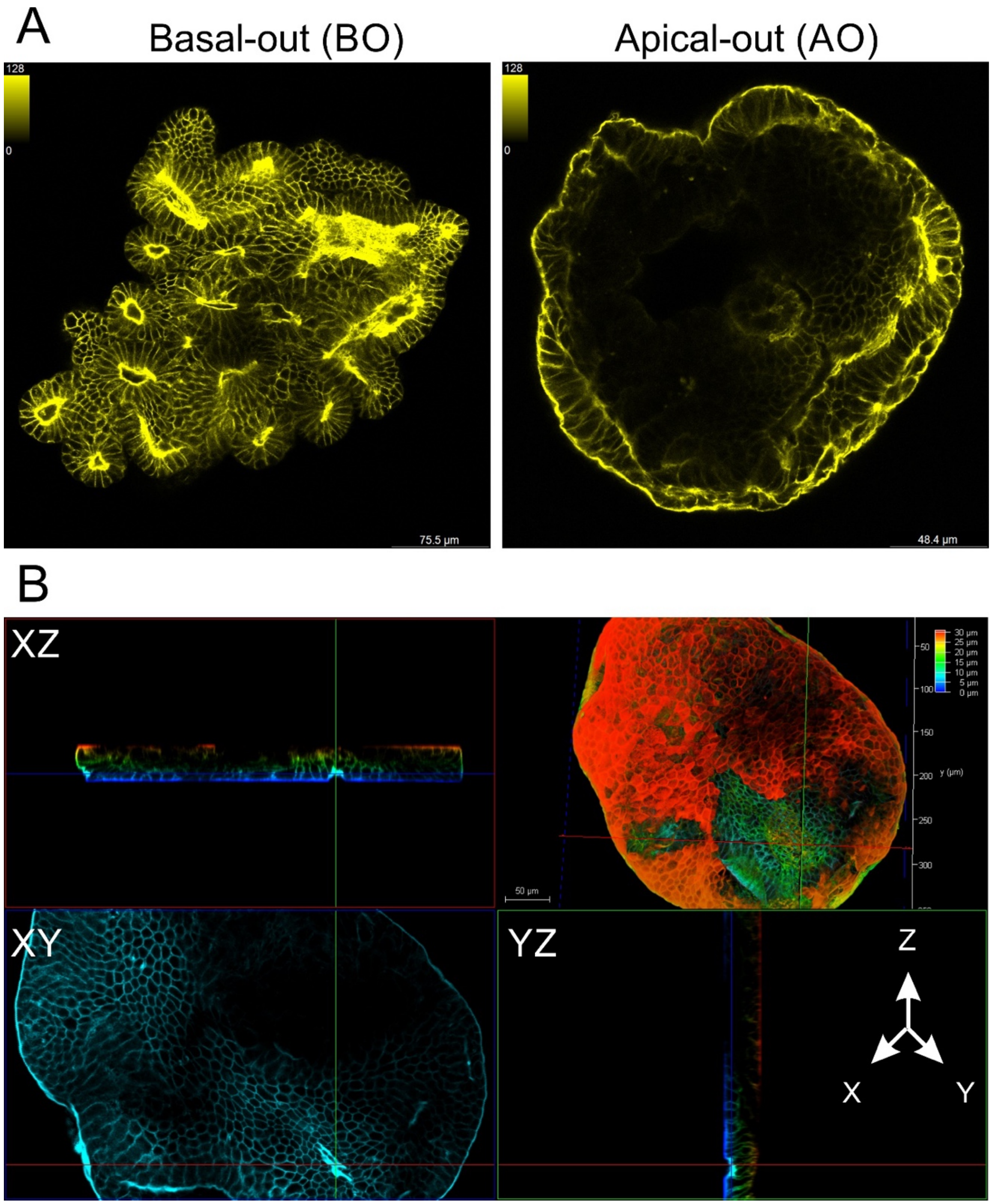


**Figure S4. Representative confocal fluorescence microscopy images of polarity reverted (AO) and BO organoids, with F-actin labeling of apical membrane with phalloidin-Alexa 546 conjugate. A:** XY optical sections of BO (Matrigel embedded organoid) and AO (1 day old suspension of organoids on a low attachment surface) pig intestinal organoids. **B:** Depth color-coded 3D reconstruction of AO organoid, imaged in glass mounted sample. Scale bars are indicated.

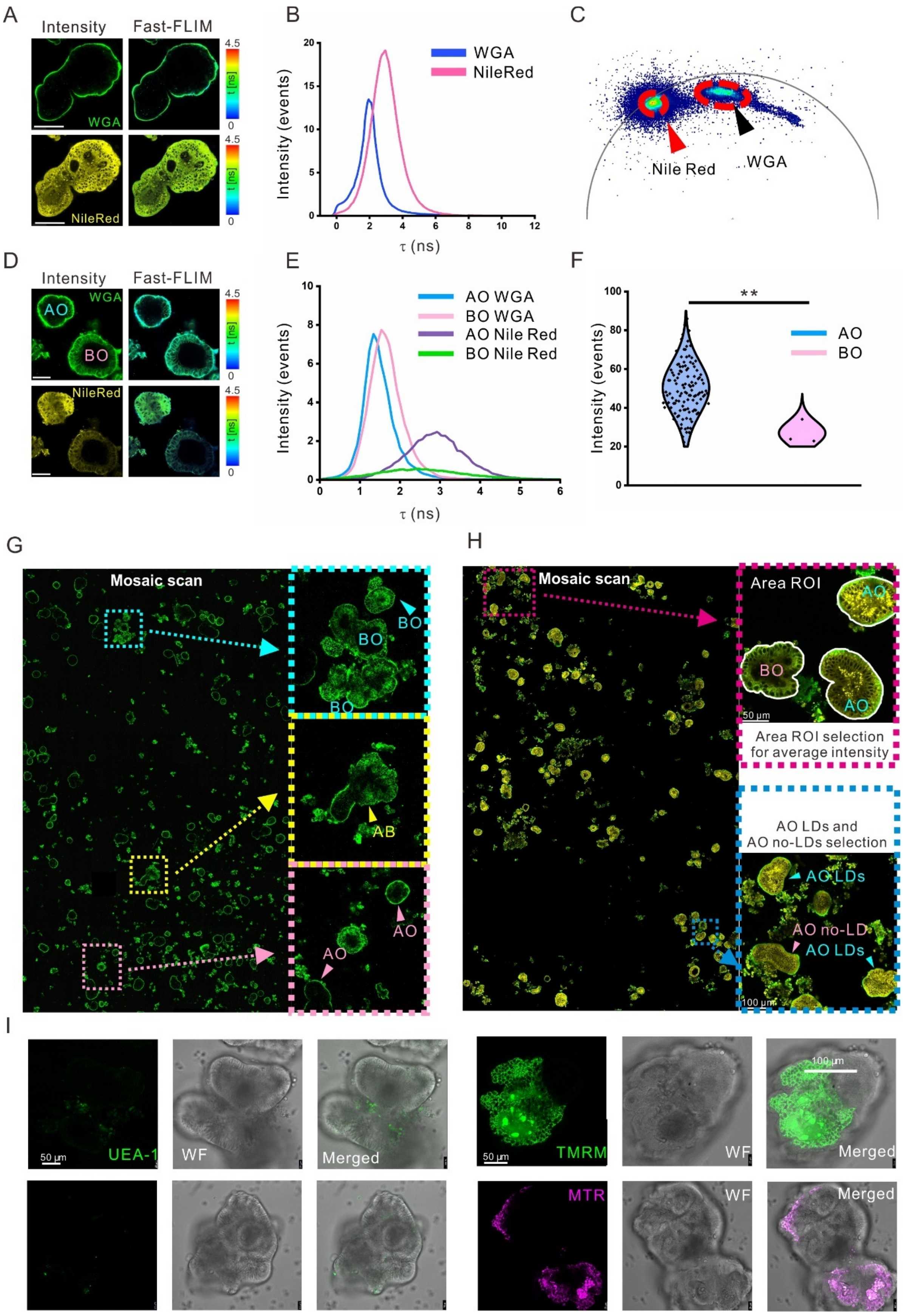

A
Intensity
Fast-FLIM
WGA
NileRed
4.5
0
B
WGA
NileRed
Intensity (events)
τ (ns)
C
Nile Red
WGA
D
Intensity
Fast-FLIM
AO
BO
WGA
NileRed
E
AO WGA
BO WGA
AO Nile Red
BO Nile Red
Intensity (events)
τ (ns)
F
**
AO
BO
Intensity (events)
G
Mosaic scan
BO
AB
AO
H
Mosaic scan
Area ROI
AO
BO
50 μm
Area ROI selection for average intensity
AO LDs and AO no-LDs selection
AO LDs
AO no-LDs
100 μm
I
UEA-1
WF
Merged
50 μm
TMRM
MTR
100 μm

**Figure S5. Evaluation of live organoid staining with WGA-Alexa Fluor 488, Nile Red and other live imaging dyes. A:** Representative confocal fluorescence intensity and fast-FLIM images of an apical-out porcine small intestinal organoid labeled with WGA and Nile Red. Scale bar is 100 μm. **B, C:** Fluorescence lifetime distribution histograms and phasor plot for WGA and Nile Red from the apical-out organoid shown in A (WGA ~2.1 ns, Nile Red ~3.3 ns). **D, E:** Comparison of WGA and Nile Red fluorescence intensity (D) and normalized lifetime distribution (E) in AO and BO organoids. Scale bar is 50 μm. **F:** Violin plots display average Nile Red fluorescence intensity in AO and BO organoids (AO area ROI counts: 119, BO area ROI counts: 3, intensity measured in FIJI software, p=0.006). **G:** WGA staining identifies three types of small intestinal organoids. Left: large mosaic scan of organoids in a single well, *Right:* representative images of BO, AB and AO from the mosaic scan. **H:** Analysis of WGA and Nile Red co-stained organoids. *Left:* Mosaic scan image. *Top right*: Organoid area ROI selection for average Nile Red intensity measurement using freehand tool in FIJI software. *Bottom right:* Counting the number of AO with lipid droplets (AO LDs) and without lipid droplets (AO no-LDs). **I:** Representative images of staining of polarity-reverted small intestinal organoids with UEA-1, MitoTracker Green, TMRM and MitoTracker Red.

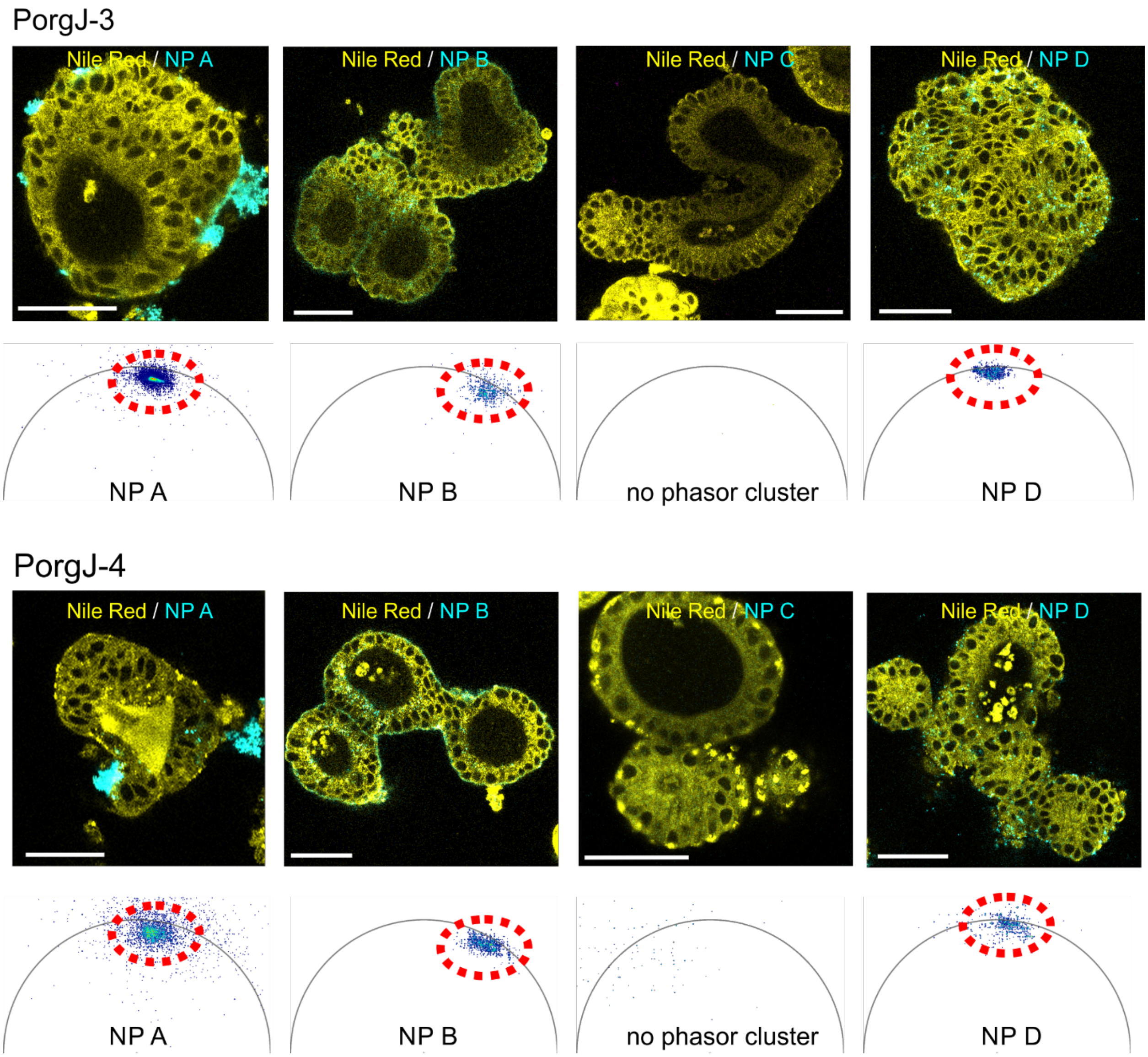

**Figure S6**: **Reproducibility of MNP uptake patterns across independent porcine intestinal organoid lines.** Two additional independent porcine intestinal organoid lines (PorgJ-3 and PorgJ-4) were exposed to 4 NP types (A-D, shown in blue) and co-stained with Nile Red (shown in yellow). Both lines displayed similar uptake patterns to PorgJ-2: NP A displayed apical membrane accumulation, NP B and NP D displayed intracellular uptake, confirmed with phasor FLIM analysis. Scale bar is 50 µm.

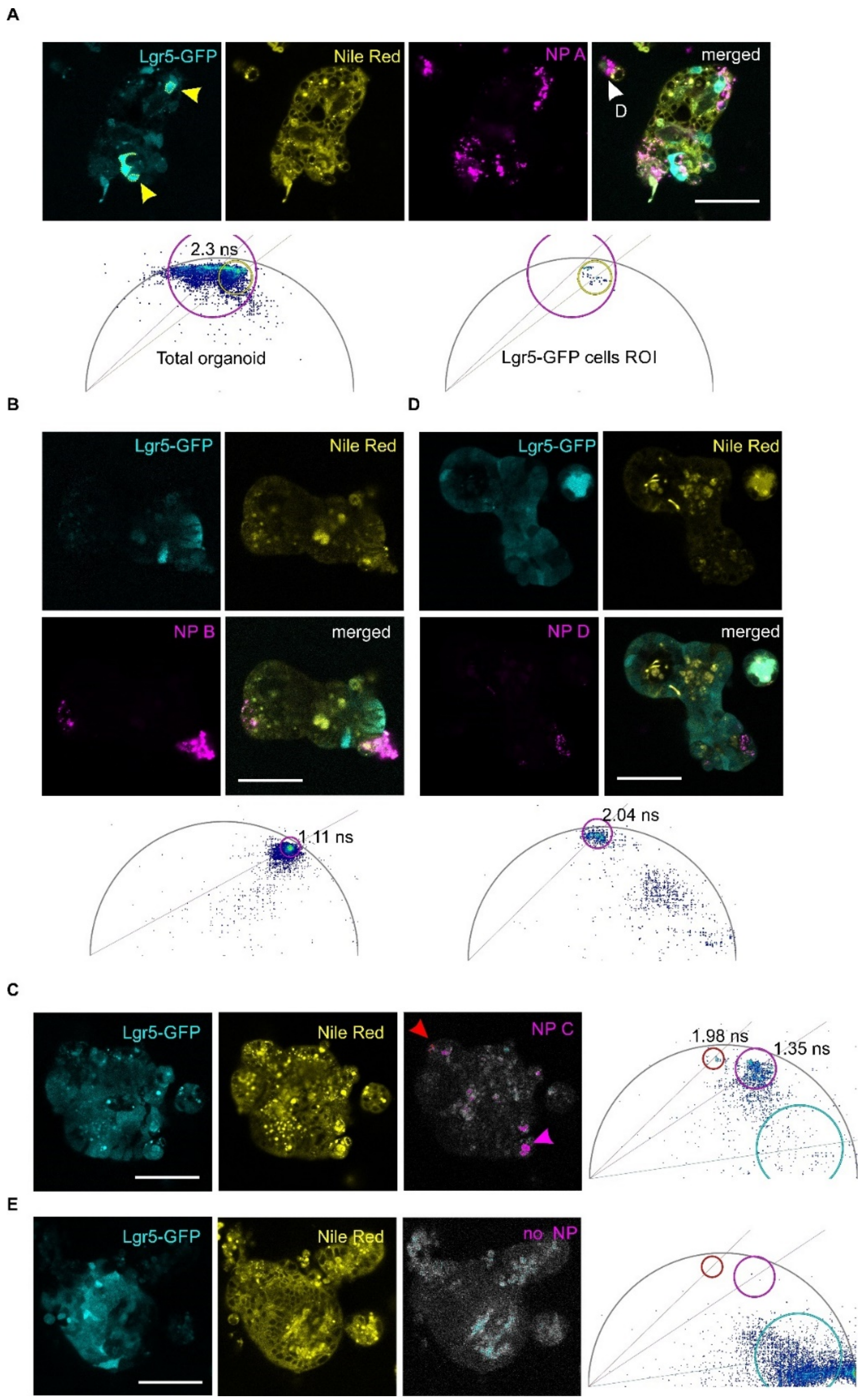


**Figure S7**. **Analysis of MNP uptake in polarity-reverted mouse Lgr5-GFP small intestinal organoids.** Organoids were polarity-reverted and incubated with NP A-D, essentially as with pig intestinal organoids (see Methods section). Lgr5-GFP organoids display expression of Lgr5-GFP (stem cell marker, shown in cyan), which disappears in differentiated cells (partially co-stained with Nile Red, yellow). NPs are shown in magenta. **A:** Local NP A uptake by specialized cell clusters, representative GFP negative or weak staining in organoids. NP A signal was also detected in cell debris, shown with white arrow (D). Yellow arrows indicate highly intense Lgr5-GFP positive cells chosen for ROI-based phasor analysis (yellow circular ROI on the corresponding phasor plot). Magenta circular ROI on the phasor plot shows overall NP A cluster. **B, C, D:** NP types B, C and D uptake was observed only in specialized cell clusters of differentiated cells. **C:** NP type C uptake was detected by appearance of unique phasor clusters (red and magenta circular ROI on the phasor plot correspond to arrows and red and magenta color mask on the NP C intensity image) with characteristic fluorescence lifetime ($\tau_\phi$ range of 1.98-1.35 ns). **E:** Unstained control group organoids did not display fluorescence with the characteristic lifetime of NP C (no phasor clusters in the red and magenta circular ROIs on the phasor plot). Cyan circular phasor ROI corresponds to intrinsic fluorescence noise signal of organoids in the spectral channel of dibutoxy-aza-BODIPY dye. Phasor plot was made as a sum of 8 phasor plots from corresponding FLIM microscopy data of individual organoids from the control group. Phasor plots from NP A-D were reconstructed from single organoid FLIM data set (per image) in LAS X software. with threshold 7, pixel binning 1 and median filter 11. Scale bar is 50 µm.

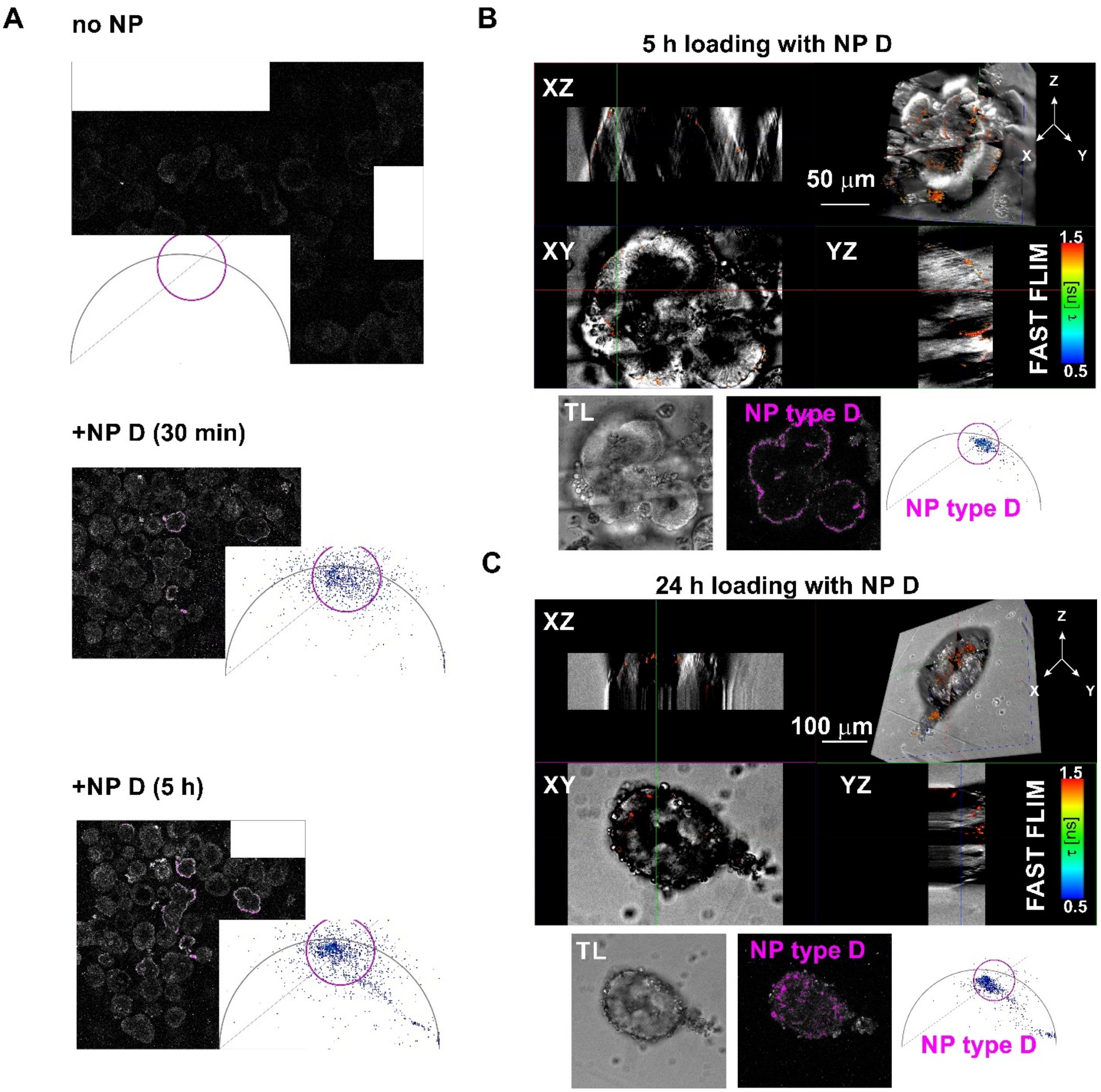

**Figure S8. Time-course analysis of NP D uptake in pig intestinal organoids**. Organoids were incubated with NP D and measured in fluorescence and transmission light channels with mosaic scanning (all the scans were performed within <15 min timeframe). **A:** Mosaic images and corresponding phasor plots of control (no NP) and NP D-treated (0.5-5 h) organoid cultures. Circular magenta phasor ROI (with $\tau_\phi$ ~2 ns) indicates the corresponding phasor cluster of NP D. **B, C:** 3D reconstruction and phasor plot of individual (middle) optical section of organoid analyzed at 5 h (B) and 24 h (C) time points. At 5 h point NP D fluorescence signal was mainly detected at the organoid periphery. At 24 h time point, NP D fluorescence signal was mainly detected inside the organoid.

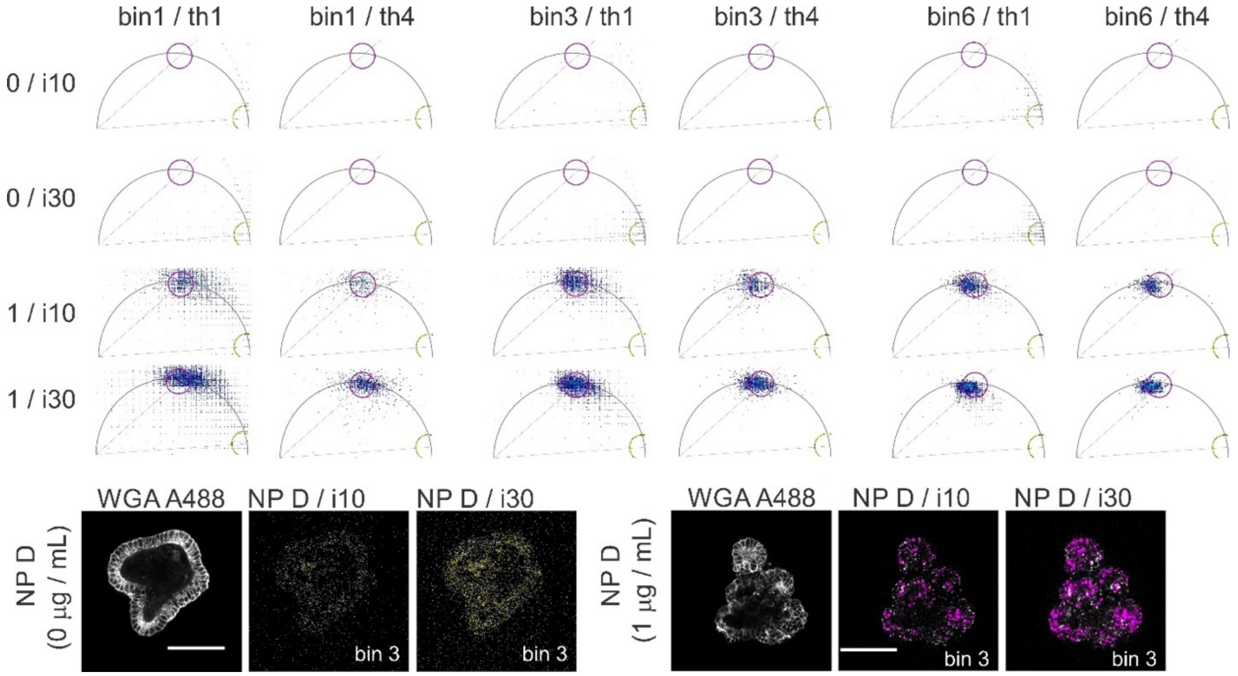


**Figure S9. Effect of excitation laser power intensity, pixel binning and threshold application on the phasor clusters of NP D in pig intestinal organoids.** Phasor plots were produced in LAS X software from unstained (referred as 0) or 1 µg / mL NP D-loaded (referred as 1) organoids images (the bottom row) acquired with either 10 % (i10) or 30 % (i30) laser power intensity. Pixel binning (bin) 1, 3 or 6 or phasor threshold (th) 1 or 4 were applied during image phasor plots reconstruction. Magenta and yellow circular phasor ROIs point at the position of the typical NP D cluster or the noise signal cluster, respectively. Yellow and magenta pseudocolor masks on the intensity images of NP D spectral channel correspond to lifetime events from the noise and NP D fluorescence signal phasor ROIs. Images were acquired with 100 Hz scanning speed, 1024x1024 resolution, pinhole 1 AU, 80 MHz laser pulse frequency, 1 frame repetition. rate. Median filter 11, harmonic 1 were applied for phasor plot reconstruction. Scale bar is 50 µm.

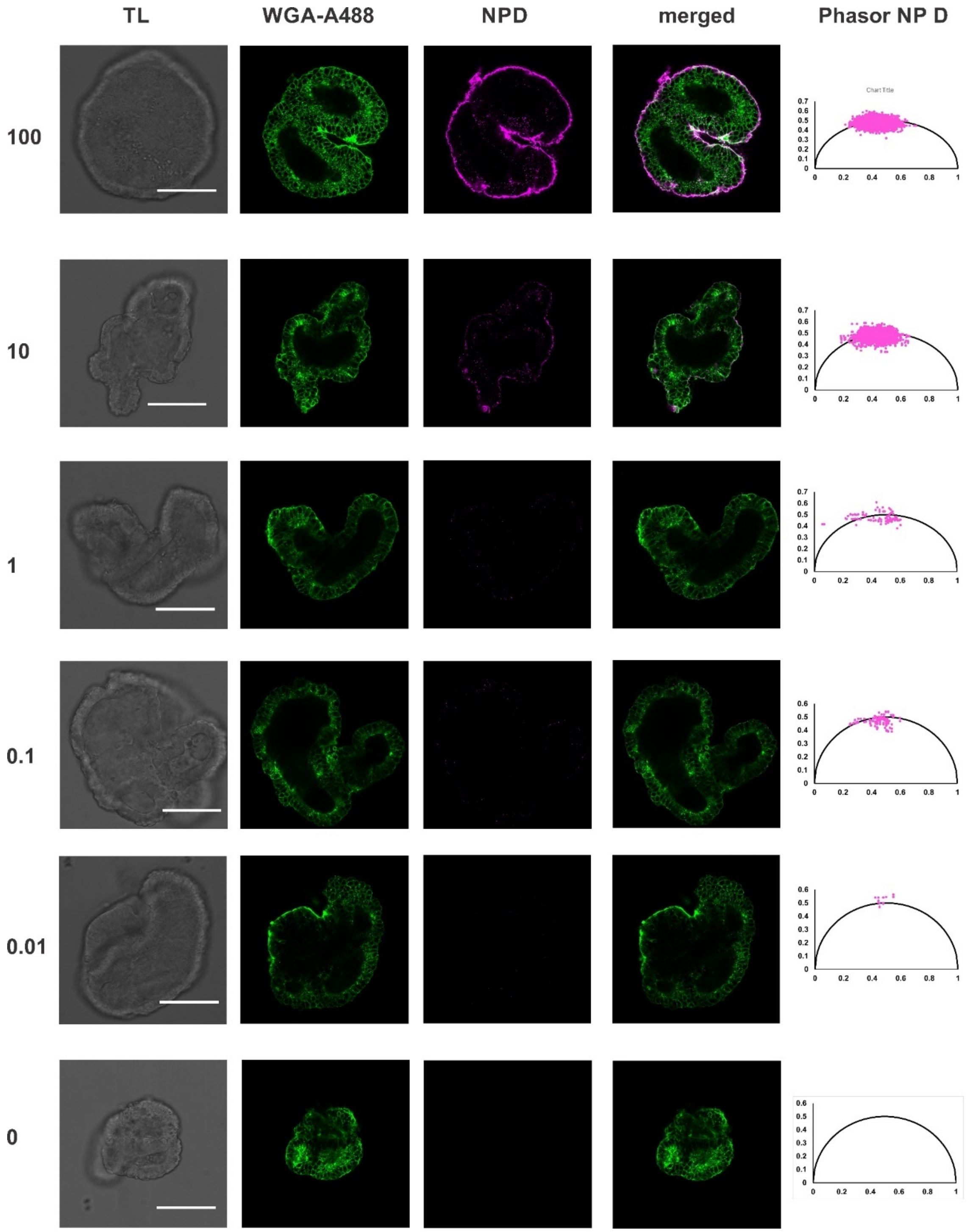


**Figure S10. Typical examples of BO pig intestinal organoids loaded with NP D at various concentrations (0, 0.01, 0.1, 1, 10 and 100 µg/mL) and co-stained with WGA-Alexa Fluor 488.** Images were acquired with pixel resolution 1024x1024, frame repetition 1, 100 Hz, pinhole 1AU, 80 MHz pulse repetition rate. Scale bar is 50 µm. Corresponding phasor plots were exported from phasor FLIM napari plugin as a list of G and S coordinates and reconstructed in Microsoft Excel software. The phasor plots demonstrate that the number of acquired events in NP D lifetime zone clusters decreases proportionally with the decrease in NP D loading concentration down to zero event count for unstained organoid.

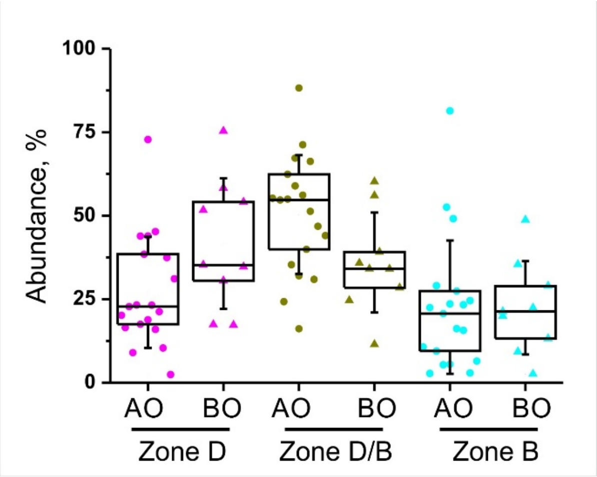


**Fig. S11. Lifetime zone analysis of phasor clusters of AO and BO organoids loaded with a mixture of D and B NP.** Boxes show 25 and 75 percentiles, while whiskers show standard deviation. Each point corresponds to individual organoid value. Mann-Whitney comparison between AO and BO organoids inside one zone group did not detect statistical difference at $p<0.05$.

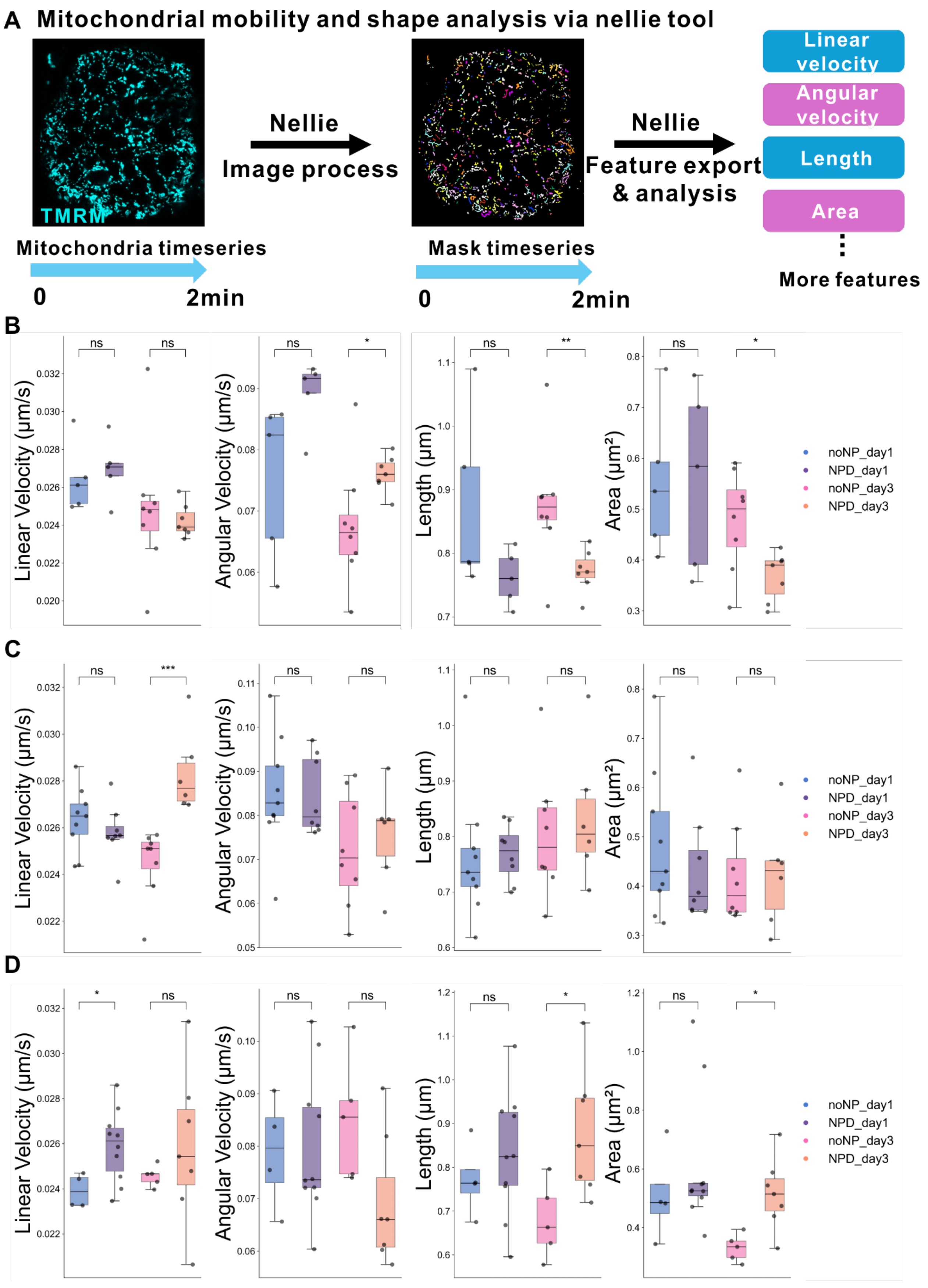

A Mitochondrial mobility and shape analysis via nellie tool
TMRM
Nellie
Image process
Nellie
Feature export & analysis
Linear velocity
Angular velocity
Length
Area
More features
Mitochondria timeseries
0
2min
Mask timeseries
0
2min
B
C
D
Linear Velocity (μm/s)
Angular Velocity (μm/s)
Length (μm)
Area (μm²)
ns
*
**
***
noNP_day1
NPD_day1
noNP_day3
NPD_day3

**Figure S12. Analysis of effects of NP D on mitochondrial morphology and dynamics in pig intestinal organoids. A:** Scheme illustrating the workflow of analysis of mitochondrial features (velocity (linear and angular), length, and area) using Nellie software, processed from from XYt time-lapse imaging (2 minutes, 47 timeframes) of TMRM fluorescence. **B, C, D:** Quantitative analysis of three independent organoid replicates (1- and 3-days exposure to NP D) showing mitochondrial features. Boxplots display average values of mitochondrial metrics per organoid (individual points represent single organoids).

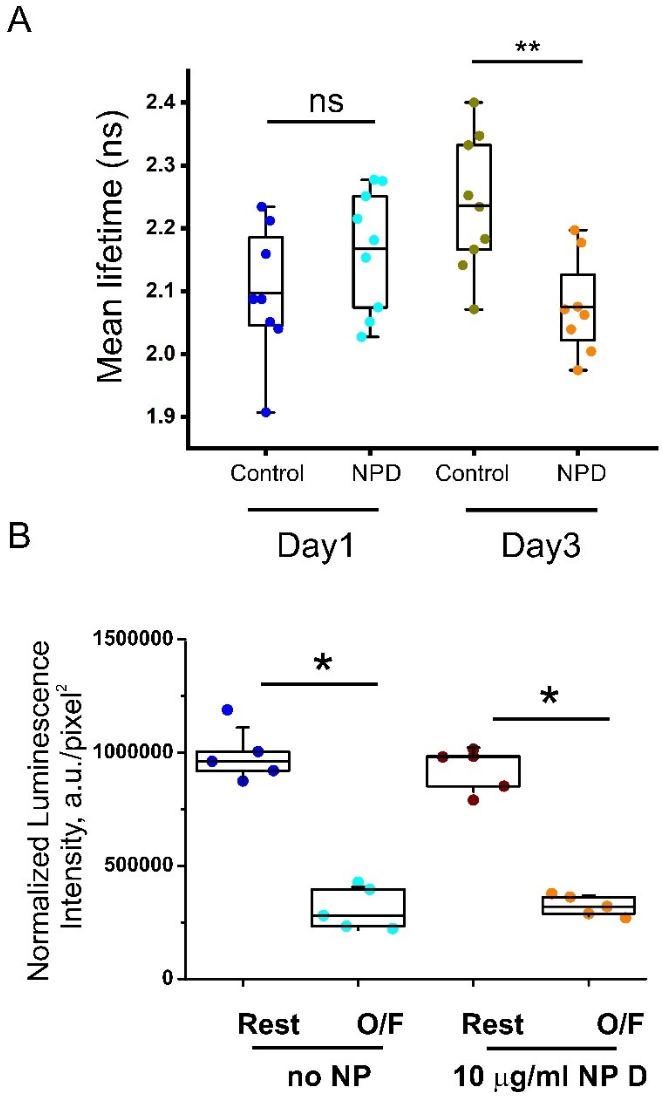


**Figure S13. Independent replicates of mitochondrial polarization analysis and ATP test in control (no NP) and NP D groups. A:** Analysis of mitochondria polarization in control and NP D loaded BO organoids by comparison of mean TMRM fluorescence lifetime. No significant difference was observed after 1-day exposure to NP D; mitochondria polarization was significantly different ($p<0.01$) after 3 days exposure to NP D. Difference in TMRM fluorescence lifetime between control and NP D-treated group was 0.16 ns. **B**: Second replicate of ATP test. No differences were observed between control and NP D treated organoids at resting (Rest) and under stimulation with Oligomycin / FCCP (O/F).

#### Supplementary tables

**Table S1. Analysis of organoid polarity phenotypes in population of organoids in suspension from 5 independent experiments on polarity reversion.** Total analysis of organoid population was done by mosaic scan imaging based on WGA-Alexa Fluor 488 conjugate staining of live organoids.

| | **Total amount of organoids analyzed** | **Percentage of AO organoids from total organoid number** | **Percentage of AB organoids from total organoid number** | **Percentage of BO organoids from total organoid number** |
|---|---|---|---|---|
| 1 | 136 | 96.3% | 0.7% | 3% |
| 2 | 156 | 94.2% | 1.9% | 3.8% |
| 3 | 102 | 98% | 0% | 2% |
| 4 | 173 | 98.8% | 0% | 1.2% |
| 5 | 98 | 98% | 1% | 1% |

**Table S2.** The number and percentage of AO, partial AO and BO observed in 4 mosaic scan images, calculated based on WGA and F-actin staining patterns separately, as shown on Figures 2C and S3.

| | | **Numbers of AO** | **Numbers of Partial AO** | **Numbers of BO** | **Total** | **Percentage of AO/All organoid** | **Percentage of partial AO/All organoid** | **Percentage of BO/All organoid** | **Note** |
|---|---|---|---|---|---|---|---|---|---|
| 1 | WGA | 60 | 5 | 0 | 65 | 92.3% | 7.7% | 0% | 1 F-actin staining organoid without WGA |
| | F-actin | 63 | 3 | 0 | 66 | 95.4% | 4.5% | 0% | |
| 2 | WGA | 79 | 7 | 1 | 87 | 90.9% | 8% | 1.1% | 1 F-actin |

| | | | | | | | | | |
|---|---|---|---|---|---|---|---|---|---|
| | F-actin | 82 | 5 | 1 | 88 | 93.1% | 5.7% | 1.2% | staining organoid without WGA |
| 3 | WGA | 98 | 6 | 1 | 105 | 93.3% | 5.7% | 1% | |
| | F-actin | 100 | 4 | 1 | 105 | 95.2% | 3.8% | 1% | |
| 4 | WGA | 116 | 5 | 0 | 121 | 95.9% | 4.1% | 0% | |
| | F-actin | 119 | 2 | 0 | 121 | 98.3% | 1.7% | 0% | |

**Table S3. Comparison of NP D uptake in AO and BO organoids over a range of concentrations (0-100 µg/mL) using fluorescence intensity and Phasor FLIM events counting approaches.** AO and BO organoid data from each concentration group were compared with Mann-Whitney test (at p = 0.05). Data represents one of two individual experimental replicates (R2), shown in Fig.4 C,D.

| **Intensity-based analysis** | | | | | | | | | |
|---|---|---|---|---|---|---|---|---|---|
| [NP D], µg / mL | N1 (AO) N2 (BO) | U | Z | Exact prob>\|U\| | Asymp. Prob >\|U\| | Median, (AO/BO) | Mean Rank (AO/BO) | Sum Rank (AO/BO) | Organoid loading events, % (AO/BO) |
| noNP | N1 = 15 N2 = 13 | 57 | -1.8426 | 0.06475 | 0.06539 | 52.1 / 64.7 | 11.8 / 17.6 | 177 / 229 | n/a |
| 0.01 | N1 = 19 N2 = 13 | 47 | -2.9161 | 0.00259 | 0.00354 | 77.8 / 91.4 | 12.5 / 22.4 | 237 / 291 | n/a |
| 0.1 | N1 = 16 N2 = 22 | 21 | -4.5679 | <0.0001 | <0.0001 | 91.5 / 177.9 | 9.8 / 26.5 | 157 / 584 | n/a |
| 1 | N1 = 17 N2 = 17 | 48 | -3.3066 | 5.56E-04 | 9.44E-04 | 59.7 / 155.7 | 11.8 / 23.2 | 201 / 394 | n/a |

| | | | | | | | | | |
|---|---|---|---|---|---|---|---|---|---|
| 10 | N1 = 25 N2 = 17 | 26 | -4.7662 | <0.0001 | <0.0001 | 186.5 / 1791.7 | 14 / 32.5 | 351 / 552 | n/a |
| 25 | N1 = 23 N2 = 19 | 59 | -4.018 | <0.0001 | <0.0001 | 289.4 / 2462.8 | 14.6 / 29.9 | 335 / 568 | n/a |
| 50 | N1 = 16 N2 = 20 | 20 | -4.4411 | <0.0001 | <0.0001 | 501 / 5484.7 | 9.8 / 25.5 | 156 / 510 | n/a |
| 100 | N1 = 17 N2 = 12 | 5 | -4.2731 | <0.0001 | <0.0001 | 1653.3 / 12039.2 | 9.3 / 23.1 | 158 /277 | n/a |
| **Phasor FLIM event counting analysis** | | | | | | | | | |
| [NPD], µg / mL | N1 (AO) N2 (BO) | U | Z | Exact prob>\|U\| | Asymp. Prob >\|U\| | Median, (AO/BO) | Mean Rank (AO/BO) | Sum Rank (AO/BO) | Organoid loading events, % (AO/BO) |
| noNP | N1 = 13 N2 = 9 | 58.5 | 0 | | 1 | 0 /0 | 11.5 / 1..5 | 149.5 / 103.5 | 0 / 0 |
| 0.01 | N1 = 21 N2 = 10 | 58 | -2.8487 | 0.00508 | 0.0022 | 0 / 2.13644E-5 | 13.8 / 20.7 | 289 / 207 | 4.8 / 50 |
| 0.1 | N1 = 13 N2 = 17 | 13 | -4.0981 | <0.0001 | <0.0001 | 0 / 5.81784E-4 | 8 /21.2 | 104 / 361 | 38.5 / 100 |
| 1 | N1 = 14 N2 = 9 | 2 | -4.1128 | <0.0001 | <0.0001 | 0 / 1.2E-3 | 7.6 / 18.8 | 107 /169 | 14.3 /100 |
| 10 | N1 = 10 N2 = 7 | 0 | -3.3877 | 1.03E-04 | 7.05E-04 | 2.44E-3 / 2.508E-2 | 5.5 / 14 | 55 /98 | 60 / 100 |
| 25 | N1 = 17 N2 = 12 | 26 | -3.3436 | 3.95E-04 | 8.27E-04 | 2.07E-3 / 1.114E-2 | 10.5 /21.3 | 179 /256 | 88.2 /100 |
| 50 | N1 = 10 N2 = 11 | 0 | -3.8378 | <0.0001 | <0.0001 | 2.06E-3 / 8.34E-2 | 5.5 / 16 | 55 / 176 | 100 / 100 |
| 100 | N1 = 15 N2 = 11 | 20 | -3.2178 | 3.24E-04 | 6.46E-04 | 1.105E-2 / 0.16887 | 9.3 / 19.2 | 140 / 211 | 100 / 100 |

**Table S4**. **Comparison of NP D uptake in AO and BO organoids over a range of concentrations (0-100 µg/mL) using fluorescence intensity and Phasor FLIM events counting approaches.** AO and BO organoid data from each concentration group were compared with Mann-Whitney test (at p value 0.05). Data represents replicate R1 of 2 independent repeats.

| ***Intensity analysis*** | | | | | | | | | |
|---|---|---|---|---|---|---|---|---|---|
| [NP D], µg / mL | N1 (AO) N2 (BO) | U | Z | Exact prob>\|U\| | Asymp. Prob >\|U\| | Median, (AO/BO) | Mean Rank (AO/BO) | Sum Rank (AO/BO) | Organoid loading events, % (AO /BO) |
| noNP | N1 =23 N2 =13 | 150 | 0 | 1 | 1 | 15.774 / 16.434 | 18.5 / 18.5 | 426 /240 | n/a |
| 0.01 | N1 =26 N2 = 17 | 236 | 0.3602 | 0.72161 | 0.71871 | 13.8 / 10.02 | 22.6/21.1 | 587/359 | n/a |
| 0.1 | N1 =24 N2 = 10 | 28 | -3.4584 | 2.12E-04 | 5.43E-04 | 27.259 / 60.5275 | 13.7 / 26.7 | 328 / 267 | n/a |
| 1 | N1 =24 N2 = 27 | 99 | -4.2366 | <0.0001 | <0.0001 | 9.904 / 59.698 | 16.6 / 34.3 | 399 / 927 | n/a |
| 10 | N1 = 16 N2 = 12 | 42 | -2.4837 | 1.13E-02 | 0.013 | 160.225/833.204 | 11.1/19 | 178 / 228 | n/a |
| 25 | N1 =13 N2 = 18 | 14 | -4.1033 | <0.0001 | <0.0001 | 22.106 / 998.5075 | 8.1 / 21.7 | 105 / 391 | n/a |
| 50 | N1 = 18 N2 =16 | 78 | -2.26 | 0.0224 | 0.02382 | 5.2535 / 1922.243 | 13.8 / 21.6 | 249 / 346 | n/a |
| 100 | N1 = 13 N2 = 6 | 8 | -2.675 | 0.00472 | 0.00747 | 352.788 / 7725.37 | 7.6 / 15.2 | 99 / 91 | n/a |
| **Phasor FLIM event counting analysis** | | | | | | | | | |
| [NP D], µg / mL | N1 (AO) N2 (BO) | U | Z | Exact prob>\|U\| | Asymp. Prob >\|U\| | Median, (AO/BO) | Mean Rank (AO/BO) | Sum Rank (AO/BO) | Organoid loading events, % (AO /BO) |
| noNP | N1 =7 N2 =11 | 38.5 | 0 | 1 | 1 | 0/0 | 9.5 /9.5 | 66.5 /104.5 | 0/0 |

| | | | | | | | | | |
|---|---|---|---|---|---|---|---|---|---|
| 0.01 | N1 =12 N2 = 12 | 35 | -2.4228 | 0.01164 | 0.0154 | 0 / 3.9814E-5 | 9.4/15.6 | 113/187 | 17.7 /58.3 |
| 0.1 | N1 = 9 N2 = 9 | 0 | -3.5505 | <0.0001 | 3.85E-04 | 4.04392E-5 / 7.12172E-4 | 5//14 | 45/126 | 55.6 /100 |
| 1 | N1 = 5 N2 = 7 | 0 | -2.8104 | 2.53E-03 | 4.95E-03 | 0 / 7.31422E-4 | 3//9 | 15/63 | 20 /100 |
| 10 | N1 = 13 N2 =4 | 0 | -2.8873 | 8.40E-04 | 0.00389 | 0.00379 /0.03488 | 7/15.5 | 91/62 | 92.3 / 100 |
| 25 | N1 = 6 N2 = 8 | 14 | -1.2264 | 0.22844 | 0.22003 | 0.00965 / 0.03202 | 5.8 /8.8 | 35 /70 | 83.3 / 100 |
| 50 | N1 = 5 N2 = 9 | 2 | -2.6667 | 0.004 | 0.00766 | 0.01991 / 0.09227 | 3.4 /9.8 | 17/88 | 100 /100 |
| 100 | N1 = 5 N2 = 8 | 7 | -1.8298 | 0.06527 | 0.06728 | 0.07511 /0.23885 | 4.4/8.6 | 22/69 | 100 /100 |

**Table S5. Comparison of NP D uptake in AO organoids.** Data represents one of two individual experimental replicates (R2), shown in Fig. 4C,D.

| **Intensity-based analysis (AO organoids)** | | | | | | | |
|---|---|---|---|---|---|---|---|
| Kruskal-Wallis ANOVA | | | | Conover's post-hoc comparison to no NP group | | | |
| [NP D], µg / mL | AO median | Mean Rank | Sum Rank | Mean Rank Difference | Z | Prob | Difference at p=0.05 |
| noNP | 52.1 | 22.2 | 333 | 0 | n/a | n/a | n/a |
| 0.01 | 77.8 | 41.6 | 790 | -19.37895 | -2.26213 | 0.20184 | n.s. |
| 0.1 | 91.5 | 53.3 | 856 | -31.3 | -3.51134 | 0.006 | s.d. |
| 1 | 59.7 | 36.1 | 613 | -13.85882 | -1.57734 | 0.4839 | n.s. |
| 10 | 186.5 | 87.3 | 2182 | -65.08 | -8.0341 | 7.03E-12 | s.d. |

| | | | | | | | |
|---|---|---|---|---|---|---|---|
| 25 | 289.4 | 99.3 | 2283 | -77.06087 | -9.36173 | 4.29E-15 | s.d. |
| 50 | 501 | 110.6 | 1770 | -88.425 | -9.91983 | 1.70E-16 | s.d. |
| 100 | 1653.3 | 129.4 | 2199 | -107.15294 | -12.1956 | 2.57E-22 | s.d. |
| Test statistics (at p level 0.05): Chi-Square 100.13455, DF 7, Prob > Chi-Square is < 0.0001 | | | | | | | |
| **Phasor FLIM event counting analysis (AO organoids)** | | | | | | | |
| Kruskal-Wallis ANOVA | | | | Conover's post-hoc test comparison to no NP group | | | |
| [NP D], µg / mL | AO median | Mean Rank | Sum Rank | Mean Rank Difference | Z | Prob | Difference at p=0.05 |
| noNP | 0 | 30 | 390 | 0 | n/a | n/a | n/a |
| 0.01 | 0 | 31.5 | 661 | -1.47619 | -0.25882 | 1 | n.s. |
| 0.1 | 0 | 43.8 | 569 | -13.76923 | -2.17213 | 0.25677 | n.s. |
| 1 | 0 | 35.6 | 498 | -5.57143 | -0.89503 | 1 | n.s. |
| 10 | 2.44E-03 | 65.3 | 653 | -35.3 | -5.1928 | 1.43E-05 | s.d. |
| 25 | 2.07E-03 | 77.5 | 1317 | -47.47059 | -7.97221 | 4.23E-11 | s.d. |
| 50 | 2.06E-03 | 84.4 | 844 | -54.4 | -8.0025 | 3.80E-11 | s.d. |
| 100 | 1.11E-02 | 100.6 | 1509 | -70.6 | -11.5282 | 6.26E-19 | s.d. |
| Test statistics (at p level 0.05): Chi-Square 82.1359, DF 7, Prob > Chi-Square is < 0.0001 | | | | | | | |

**Table S6. Comparison of NP D uptake in BO organoids.** Data represents 1 of 2 individual experimental replicates (R2), shown in Fig. 4C,D.

| **Intensity-based analysis (BO organoids)** | | | | | | | |
|---|---|---|---|---|---|---|---|
| Kruskal-Wallis ANOVA | | | | Conover's post-hoc comparison to no NP group | | | |
| [NPD], µg / mL | BO median | Mean Rank | Sum Rank | Mean Rank Difference | Z | Prob | Difference at p=0.05 |
| noNP | 64.7 | 12.5 | 162 | 0 | n/a | n/a | n/a |
| 0.01 | 91.4 | 24.8 | 323 | -12.3846 | -2.16388 | 0.09713 | n.s |
| 0.1 | 177.9 | 47.4 | 1043 | -34.9476 | -6.84635 | 3.04E-09 | s.d. |
| 1 | 155.7 | 39.8 | 676 | -27.3032 | -5.07857 | 9.42E-06 | s.d. |
| 10 | 1791.7 | 81.4 | 1384 | -68.9502 | -12.8252 | 2.86E-23 | s.d. |
| 25 | 2462.8 | 85.9 | 1633 | -73.4858 | -13.9917 | 5.01E-26 | s.d. |
| 50 | 5484.7 | 109.9 | 2198 | -97.4385 | -18.7436 | 1.09E-36 | s.d. |
| 100 | 12039.2 | 124.3 | 1492 | -111.872 | -19.1516 | 1.52E-37 | s.d. |
| Test statistics (at p level 0.05): Chi-Square 114.07957, DF 7, Prob > Chi-Square is < 0.0001 | | | | | | | |
| **Phasor FLIM event counting analysis (BO organoids)** | | | | | | | |
| Kruskal-Wallis ANOVA | | | | Conover's post-hoc test comparison to no NP group | | | |
| [NPD], µg / mL | BO median | Mean Rank | Sum Rank | Mean Rank Difference | Z | Prob | Difference at p=0.05 |
| noNP | 0 | 7.5 | 67.5 | 0 | n/a | n/a | n/a |
| 0.01 | 2.14E-05 | 12.4 | 123.5 | -4.85 | -1.36381 | 0.53524 | n.s |
| 0.1 | 5.82E-04 | 31.3 | 532 | -23.7941 | -7.45755 | 1.36E-09 | s.d. |
| 1 | 1.20E-03 | 36.1 | 325 | -28.6111 | -7.84167 | 2.78E-10 | s.d. |

| | | | | | | | |
|---|---|---|---|---|---|---|---|
| 10 | 2.51E-02 | 58.6 | 410 | -51.0714 | -13.0935 | 4.63E-20 | s.d. |
| 25 | 1.11E-02 | 54.5 | 654 | -47 | -13.7711 | 3.21E-21 | s.d. |
| 50 | 8.34E-02 | 71.5 | 787 | -64.0455 | -18.4102 | 1.05E-28 | s.d. |
| 100 | 0.16887 | 76.5 | 842 | -69.0455 | -19.8475 | 8.92E-31 | s.d. |
| Test statistics (at p level 0.05): Chi-Square 77.47353, DF 7, Prob > Chi-Square is < 0.0001 | | | | | | | |

**Table S7. Comparison of NP D uptake in AO organoids (R1 independent replicate).**

| **Intensity-based analysis (AO organoids)** | | | | | | | |
|---|---|---|---|---|---|---|---|
| Kruskal-Wallis ANOVA | | | | Conover's post-hoc comparison to no NP group | | | |
| [NP D], µg / mL | AO median | Mean Rank | Sum Rank | Mean Rank Difference | Z | Prob | Difference at p=0.05 |
| 0 | 15.774 | 67.4 | 1550 | 0 | n/a | n/a | n/a |
| 0.01 | 13.802 | 70.7 | 1837 | -3.26254 | -0.26517 | 1 | n.s. |
| 0.1 | 27.259 | 75.8 | 1818 | -8.3587 | -0.66645 | 1 | n.s. |
| 1 | 9.904 | 61.3 | 1471 | 6.09964 | 0.48633 | 1 | n.s. |
| 10 | 160.225 | 109.4 | 1751 | -42.0462 | -3.00488 | 7.48E-02 | n.s. |
| 25 | 22.106 | 82.6 | 1074 | -15.2241 | -1.02076 | 1 | n.s. |
| 50 | 5.2535 | 76.3 | 1373 | -8.88647 | -0.65697 | 1 | n.s. |
| 100 | 352.788 | 117.6 | 1529 | -50.2241 | -3.36748 | 2.51E-02 | s.d. |
| Test statistics (at p level 0.05): Chi-Square 22.8339, DF 7, Prob > Chi-Square is 0.00182 | | | | | | | |
| **Phasor FLIM event counting analysis (AO organoids)** | | | | | | | |
| Kruskal-Wallis ANOVA | | | | Conover's post-hoc test comparison to no NP group | | | |
| [NP D], µg / mL | AO median | Mean Rank | Sum Rank | Mean Rank Difference | Z | Prob | Difference at p=0.05 |
| 0 | 0 | 14 | 98 | 0 | n/a | n/a | n/a |
| 0.01 | 0 | 16.4 | 197 | -2.41667 | -0.55608 | 1 | n.s. |
| 0.1 | 4.E-05 | 24.4 | 220 | -10.4444 | -2.26805 | 0.27349 | n.s. |

| | | | | | | | |
|---|---|---|---|---|---|---|---|
| 1 | 0 | 17.6 | 88 | -3.6 | -0.67283 | 1 | n.s. |
| 10 | 3.79E-03 | 43.1 | 560 | -29.0769 | -6.78752 | 2.10E-07 | s.d. |
| 25 | 9.65E-03 | 42.3 | 254 | -28.3333 | -5.57324 | 1.47E-05 | s.d. |
| 50 | 1.99E-02 | 50.6 | 253 | -36.6 | -6.8404 | 1.80E-07 | s.d. |
| 100 | 7.51E-02 | 56.6 | 283 | -42.6 | -7.96178 | 3.07E-09 | s.d. |
| Test statistics (at p level 0.05): Chi-Square 45.90195, DF 7, Prob > Chi-Square is < 0.0001 | | | | | | | |

#### Table S8. Comparison of NP D uptake in BO organoids (R1 independent replicate).

| **Intensity-based analysis (BO organoids)** | | | | | | | |
|---|---|---|---|---|---|---|---|
| Kruskal-Wallis ANOVA | | | | Conover's post-hoc comparison to no NP group | | | |
| [NP D], µg / mL | BO median | Mean Rank | Sum Rank | Mean Rank Difference | Z | Prob | Difference at p=0.05 |
| 0 | 16.434 | 20.2 | 262 | 0 | n/a | n/a | n/a |
| 0.01 | 10.024 | 23 | 391 | -2.84615 | -0.4573 | 0.64835 | s.d. |
| 0.1 | 60.5275 | 54.4 | 544 | -34.2462 | -4.81974 | 6.43E-05 | s.d. |
| 1 | 59.698 | 43.9 | 1186 | -23.7721 | -4.16864 | 6.70E-04 | s.d. |
| 10 | 833.204 | 79.5 | 954 | -59.3462 | -8.77584 | 4.40E-13 | s.d. |
| 25 | 998.5075 | 85.8 | 1545 | -65.6795 | -10.6822 | 2.20E-17 | s.d. |
| 50 | 1922.243 | 98.7 | 1577 | -78.4087 | -12.4309 | 2.52E-21 | s.d. |
| 100 | 7725.37 | 113.5 | 681 | -93.3462 | -11.1962 | 1.56E-18 | s.d. |
| Test statistics (at p level 0.05): Chi-Square 91.38245, DF 7, Prob > Chi-Square is < 0.0001 | | | | | | | |
| **Phasor FLIM event counting analysis (BO organoids)** | | | | | | | |
| Kruskal-Wallis ANOVA | | | | Conover's post-hoc test comparison to no NP group | | | |
| [NP D], µg / mL | BO median | Mean Rank | Sum Rank | Mean Rank Difference | Z | Prob | Difference at p=0.05 |
| 0 | 0 | 8.5 | 93.5 | 0 | n/a | n/a | n/a |
| 0.01 | 3.98E-05 | 15.2 | 182.5 | -6.70833 | -3.63999 | 0.00228 | s.d. |
| 0.1 | 7.12E-04 | 30.7 | 276 | -22.1667 | -11.1703 | 4.39E-15 | s.d. |
| 1 | 7.31E-04 | 32.6 | 228 | -24.0714 | -11.2765 | 3.17E-15 | s.d. |
| 10 | 3.49E-02 | 47.3 | 189 | -38.75 | -15.032 | 1.09E-20 | s.d. |
| 25 | 3.20E-02 | 45.6 | 365 | -37.125 | -18.0965 | 1.33E-24 | s.d. |
| 50 | 9.23E-02 | 55.3 | 498 | -46.8333 | -23.6005 | 1.21E-30 | s.d. |

| | | | | | | | |
|---|---|---|---|---|---|---|---|
| 100 | 0.23885 | 64.3 | 514 | -55.75 | -27.1752 | 5.64E-34 | s.d. |
| Test statistics (at p level 0.05): Chi-Square 63.96944, DF 7, Prob > Chi-Square is < 0.0001 | | | | | | | |

**Table S9. Cluster point classification analysis of organoids loaded with a mixture of D and B NPs.**

| Organoid number | Polarity topology | Percentage of points in cluster per zone | | |
|---|---|---|---|---|
| | | G ≤ 0.5275 (zone D) | 0.5275<G<0.6532 ('D+B' zone) | G ≥ 0.6532 (zone B) |
| B_D_1 | BO | 58.3048919 | 39.13538111 | 2.55972696 |
| B_D_2_1 | AO | 23.2179226 | 71.28309572 | 5.49898167 |
| B_D_3_1 | AO | 38.4615385 | 56.1965812 | 5.34188034 |
| B_D_4_2 | AO | 18.8425303 | 32.03230148 | 49.1251682 |
| B_D_7_1 | AO | 17.481203 | 58.92857143 | 23.5902256 |
| B_D_7_2 | AO | 43.9655172 | 35.34482759 | 20.6896552 |
| B_D_8 | AO | 21.2536729 | 51.3222331 | 27.424094 |
| B_D_9 | AO | 15.9645233 | 54.98891353 | 29.0465632 |
| B_D_10 | AO | 43.902439 | 39.9113082 | 16.1862528 |
| B_D_11_1 | AO | 23.2502966 | 67.25978648 | 9.48991696 |
| B_D_11_2 | AO | 37.4620829 | 46.86552073 | 15.6723964 |
| B_D_11_3 | AO | 8.96551724 | 88.27586207 | 2.75862069 |
| B_D_12_1 | AO | 72.8323699 | 24.27745665 | 2.89017341 |
| B_D_12_2 | AO | 10.3862661 | 66.26609442 | 23.3476395 |
| B_D_12_3 | AO | 22.7777778 | 54.72222222 | 22.5 |
| B_D_13 | AO | 16.4961637 | 30.94629156 | 52.5575448 |
| B_D_14_1 | AO | 45.2599388 | 44.03669725 | 10.7033639 |
| B_D_14_2 | AO | 2.42460083 | 16.14429332 | 81.4311059 |
| B_D_14_3 | AO | 31.1151079 | 62.41007194 | 6.47482014 |
| B_D_15 | BO | 17.2165339 | 34.09177095 | 48.6916951 |
| B_D_16 | AO | 20.1932367 | 55.24154589 | 24.5652174 |
| B_D_17 | BO | 17.4045802 | 60.22900763 | 22.3664122 |
| B_D_19 | BO | 30.5496829 | 34.03805497 | 35.4122622 |
| B_D_20 | ABO | 35.2546917 | 35.79088472 | 28.9544236 |
| B_D_21 | ABO | 34.7314202 | 55.99705666 | 9.27152318 |
| B_D_22 | BO | 75.3571429 | 11.42857143 | 13.2142857 |
| B_D_23 | BO | 51.6393443 | 28.41530055 | 19.9453552 |
| B_D_24 | BO | 54.0669856 | 24.64114833 | 21.291866 |

**Table S10. The list of intestinal organoid lines used in the work.**

| Name | Species | Sex | Age | Date of primary culture isolation | Special features | Experiment type |
|---|---|---|---|---|---|---|
| mSIO Lgr5GFP | Mus musculus | female | N/A | 2009 | Lgr5-GFP labeled stem cells, from duodenum + jejunum | NP uptake test |
| porgj_2 | Sus scrofa domesticus | female | 7 weeks | 11/24/2022 | small intestinal organoids from pig jejunum | NP uptake test; ATP test, mitochondria analysis; chemokine expression |
| porgj_3 | Sus scrofa domesticus | female | 6-7 weeks | 06/_/2024 | small intestinal organoids from pig jejunum | NP uptake test; MNP co-loading test; phasor event method validation; chemokine expression |
| porgj_4 | Sus scrofa domesticus | female | 6-7 weeks | 06/_/2024 | small intestinal organoids from pig jejunum | NP uptake test |
| PigD | Sus scrofa domesticus | female | 3 weeks | 6/1/2024 | small intestinal organoids from pig jejunum | chemokine expression |
| Pig13 | Sus scrofa domesticus | male | 5 weeks | 7/1/2024 | small intestinal organoids from pig jejunum | chemokine expression |
| Pig24 | Sus scrofa domesticus | female | 6 weeks | 7/1/2024 | small intestinal organoids from pig jejunum | chemokine expression |

#### Supplementary videos (uploaded separately)

**Supplementary Video S1**: Time-lapse imaging (2 min) of NP D-exposed basal-out organoid (BO) after 3 days of treatment, labeled with WGA (green), TMRM (yellow), and NP D (magenta).

**Supplementary Video S2:** Time-lapse imaging (2 min) of control basal-out organoid (BO) after 3 days without NPD treatment, labeled with WGA (green) and TMRM (yellow).

**Supplementary Video S3:** Instruction for G, S coordinate extraction from raw .PTU files using custom Python code based on napari-flim-phasor-plotter plugin.

**Supplementary Video S4:** Time-lapse TMRM-FLIM (2 min) showing mitochondrial dynamics in organoids. FLIM data displayed with lifetime range 0-4 ns. Scale bar: 10 µm.
